# Adipocyte vesicles: ‘all-in-one’ packages that stimulate tumor mitochondrial metabolism and dynamics

**DOI:** 10.1101/649327

**Authors:** Emily Clement, Ikrame Lazar, Camille Attané, Lorry Carrié, Stéphanie Dauvillier, Manuelle Ducoux-Petit, Thomas Menneteau, Mohamed Moutahir, Sophie Le Gonidec, Stéphane Dalle, Philippe Valet, Odile Burlet-Schiltz, Catherine Muller, Laurence Nieto

**Affiliations:** Institut de Pharmacologie et de Biologie Structurale (IPBS), Université de Toulouse, CNRS, UPS, 31077 Toulouse, France; Institut des Maladies Métaboliques et Cardiovasculaires (I2MC), Université de Toulouse, INSERM, UPS, 131432 Toulouse, France; Department of Dermatology, Centre Hospitalier Lyon Sud, 69310 Pierre Bénite Cedex, France

**Keywords:** exosome, fatty acid oxidation, lipophagy, melanoma, obesity

## Abstract

Extracellular vesicles are emerging key actors in adipocyte communication. Notably, small extracellular vesicles shed by adipocytes promote melanoma aggressiveness through fatty acid oxidation, with a heightened effect in obesity. However, the vesicular actors and cellular processes involved remain largely unknown. Here, we elucidate the mechanisms linking adipocyte extracellular vesicles to metabolic remodeling and cell migration. We show that adipocyte vesicles stimulate melanoma fatty acid oxidation by providing both enzymes and substrates. In obesity, the heightened effect of extracellular vesicles depends on increased transport of fatty acids, not fatty acid oxidation-related enzymes. These fatty acids, stored within lipid droplets in cancer cells, drive fatty acid oxidation after release through lipophagy. This increase in mitochondrial activity redistributes mitochondria to membrane protrusions of migrating cells, which is necessary to increase cell migration in the presence of adipocyte vesicles. Our results provide key insights into the role of extracellular vesicles in the metabolic cooperation that takes place between adipocytes and tumors with particular relevance in obesity.

## Introduction

As worldwide obesity rates continue to climb, excess body fat has emerged as a major public health issue, given its associated complications such as cardiovascular diseases, diabetes and cancer (De Pergola & Silvestris, 2013; Park et al, 2014). Obesity is a recognized factor to increase cancer incidence and progression (Gallagher & LeRoith, 2015). This association occurs in melanoma, a skin cancer that develops from transformed melanocytes, pigment-producing cells that reside at the junction between the epidermis and the dermis. Given their high propensity to invade adjacent tissues, including subcutaneous adipose tissue (AT), and to metastasize to distant organs, melanoma is the most aggressive cutaneous cancer. Studies in murine melanoma models and epidemiological data indicate obesity is an established risk factor for melanoma incidence and progression [for a recent review, see (Clement et al, 2017)]. Adipocytes, the main component of the AT, reside in many cancer microenvironments and contribute to tumor progression through soluble factors, such as leptin or interleukin 6, and extracellular matrix remodeling (Andarawewa et al, 2005; Dirat et al, 2011; Duong et al, 2017). Recent findings highlight a metabolic cooperation between adipocytes and tumor cells, which is proving to be a key process in their tumor-promoting effects (Balaban et al, 2017; Nieman et al, 2011; Wang et al, 2017; Zhang et al, 2018).

Cellular components in the tumor microenvironment are major regulators of tumor metabolism, driving cancer cells to favor certain metabolic pathways (Gouirand et al, 2018). In particular, adipocytes provide a local supply of fatty acids (FA) that can serve as an energy source for tumors. Indeed, tumor secretions trigger lipolysis in neighboring adipocytes, and the released FA fuel fatty acid oxidation (FAO) in tumors, which increases tumor aggressiveness (Balaban et al, 2017; Nieman et al, 2011; Wang et al, 2017; Zhang et al, 2018). FAO has recently emerged as a pro-tumoral pathway involved in cancer cell proliferation, stemness and invasion (Carracedo et al, 2013; Kuo & Ann, 2018), but the molecular mechanisms behind this effect remain elusive.

Even though adipocytes have a heightened effect on tumor progression in obesity (Duong et al, 2017; Nieman et al, 2013), we still do not understand whether their metabolic cooperation with tumor cells is responsible. Until now, studies showed that metabolic exchanges between adipocytes and tumor cells requires a close proximity between the two cells types in order for tumors to provoke adipocyte lipolysis. For most types of cancer, this process can only occur at the later stages of cancer progression, when tumors become invasive and penetrate local AT or metastasize to adipocyte-rich environments (Laurent et al, 2019; Nieman et al, 2011; Wang et al, 2017; Zhang et al, 2018). However, whether adipocytes could also influence tumors at distance, for example during the early stages of disease progression before cancer cells infiltrate surrounding adipose tissue, remains elusive. We predict that extracellular vesicles (EV) are key contributors in such a process. They allow the transfer of biomolecules, including nucleic acids, proteins and lipids, to distant cells, since they diffuse through tissues and circulate in body fluids (Shah et al, 2018). Previously, we demonstrated that adipocyte EV are key participants in melanoma progression (Lazar et al, 2016). Indeed, adipocytes secrete great quantities of EV that specifically carry proteins involved in lipid metabolism, including FAO enzymes. Melanoma cells internalize these EV, and this increases FAO, which promotes tumor aggressiveness. In obesity, adipocytes secrete more EV and, when used at equal concentrations, these EV elicit a heightened effect on melanoma migration, indicating qualitative changes in EV cargo. We predict this changes likely involves FAO actors, which remain to be determined. Although the increase in tumor aggressiveness in response to adipocyte EV is dependent on FAO, the cellular processes that link this metabolic remodeling to cell migration remains unknown.

Our results reveal a mechanism by which naive adipocytes (unaltered by tumor cells) influence tumor metabolism. EV secreted by these adipocytes transfer both protein machinery and FA substrates required for FAO. We further show that the heightened effect of adipocyte EV in obesity depends on increased FA levels, whereas FAO-related protein levels remain unchanged. Cytoplasmic lipid droplets store FA transferred from adipocytes to melanoma cells by EV and release these lipids by lipophagy to fuel FAO. Finally, we propose a mechanism that links adipocyte-induced FAO to tumor cell migration, mitochondrial dynamics.

## Results

### Adipocyte EV transfer proteins involved in FA metabolism to melanoma cells

Although we know that adipocyte EV are enriched in proteins involved in fatty acid metabolism and that they increase melanoma migration through a process dependent on FAO (Lazar et al, 2016), the link remains correlative. So, we developed an experimental workflow using SILAC (Stable Isotope Labeling of Amino Acids in Cell Culture) to perform an unsupervised analysis of adipocyte proteins transferred to recipient tumor cells by EV (Fig 1A). We identified 2111-labeled proteins in adipocyte EV (Supplementary Table 1), which equates to approximately 85% of all proteins identified in EV (Fig 1B). In melanoma cells exposed to these EV, we detected 587 proteins containing heavy amino acids, which indicates they were transferred from adipocytes. In cells not exposed to EV, we only aberrantly detected 5 ‘labeled’ proteins, which underlines the robustness of this technique. Consequently, we eliminated these proteins from the list of labeled proteins. Our results indicated that approximately 30% of adipocyte EV proteins are effectively transferred to melanoma cells. Many abundant proteins within EV, such as FASN, were not transferred to melanoma, indicating a selective transfer and/or uptake of material. Amongst the proteins effectively transferred from adipocytes to melanoma cells by EV, we identified mitochondrial FAO enzymes (Fig 1C). We also identified proteins involved in FA transport and storage, as well as oxidative phosphorylation (Supplementary Table 1). Thus, using an adapted SILAC technique, we conclude that FAO-related proteins are not only present in adipocyte EV, but also efficiently transferred to melanoma cells.

**Figure 1:**
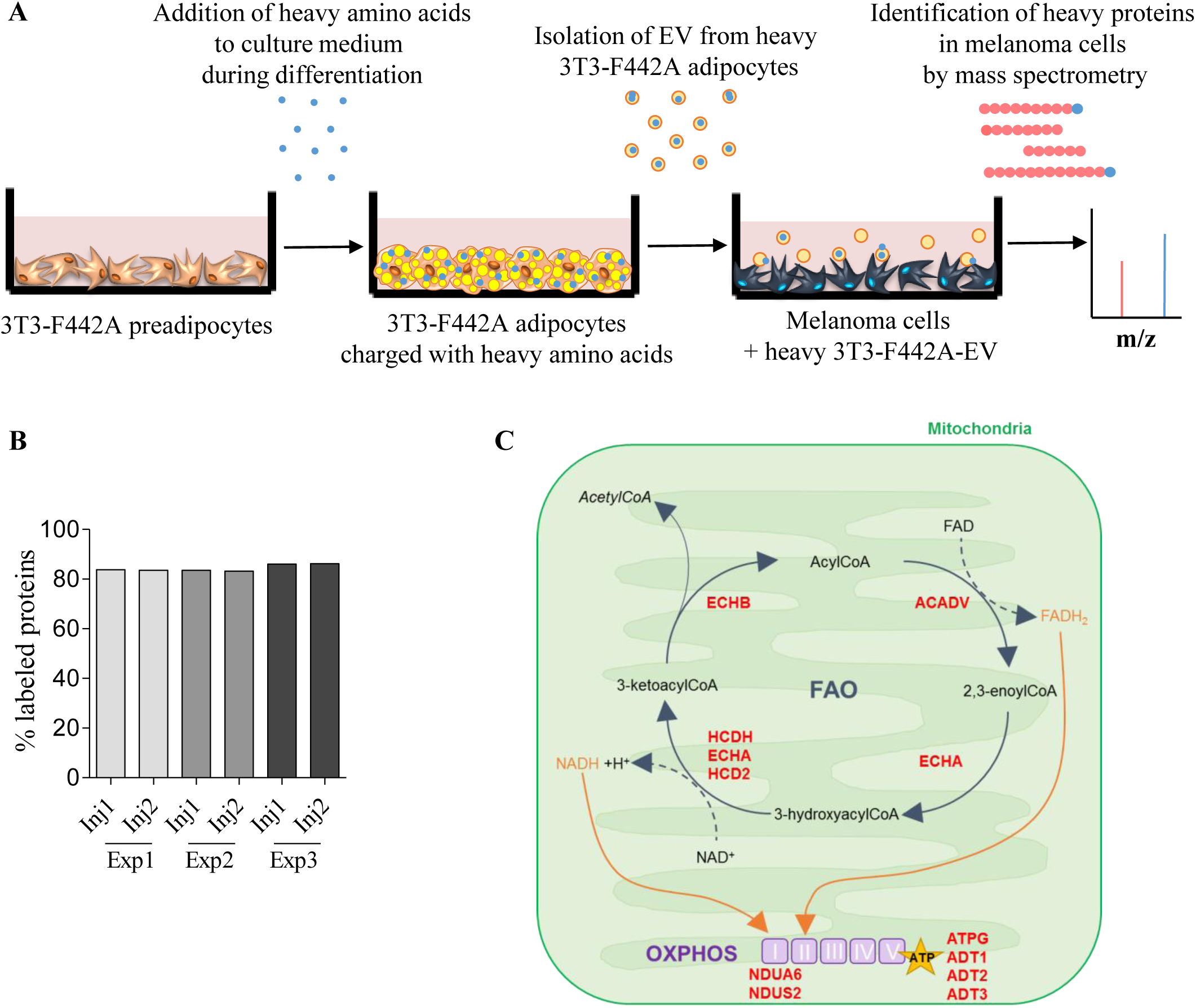
Adipocyte EV transfer proteins involved in FA metabolism to melanoma cells. A) Workflow of the SILAC approach. 3T3-F442A cells were seeded and differentiated in the presence of heavy amino acids. After 14 days of differentiation, the EV secreted by the mature labeled adipocytes were isolated and analyzed by mass spectrometry to evaluate the presence of heavy amino acid-containing proteins. These EV were also added to SKMEL28 cells for 12h, then LC-MS/MS analysis was performed to identify heavy amino-acid containing proteins that had been transferred from adipocytes to melanoma cells *via* EV. B) Three independent samples (Exp 1 to 3) of EV secreted by labeled 3T3-F442A cells were analyzed by mass spectrometry (in duplicate injections, Inj1/2). The percentage of proteins bearing at least one peptide containing a heavy amino acid is indicated. C) Proteins involved in FAO and oxidative phosphorylation (OXPHOS) that are transferred from adipocytes to melanoma cells *via* EV are shown in red.

### The heightened effect of adipocyte EV in obesity does not depend on increased FAO protein transfer

FAO levels are greater in melanoma cells treated with EV from obese individuals (termed obese adipocyte EV) compared to those treated with EV secreted by lean adipocytes (Fig 2A). To decipher the underlying mechanisms responsible for increased FAO induced by obese adipocyte EV, we performed a comparative quantitative proteomics analysis of adipocyte EV from lean and obese mice (Supplementary Table 2). We identified 1557 proteins, 87 and 100 respectively more or less abundant in obese samples. We identified 20 proteins involved in mitochondrial FAO that were present in equal abundance in both samples in our proteomic analysis (Fig 2B and Supplementary Fig 1A), which we confirmed by western blot for the two key FAO enzymes, ECHA (trifunctional enzyme subunit alpha, gene name HADHA) and HCDH (Hydroxyacyl-coenzyme A dehydrogenase, gene name HADH), in murine and human samples (Fig 2C). Moreover, we found that obese adipocyte EV are not preferentially taken up by melanoma cells compared to lean EV (Supplementary Fig 1B-C). Although FAO enzyme levels increased in the presence of primary adipocyte EV, in accord with a direct transfer of these proteins by EV, as demonstrated in the SILAC experiment (Supplementary Table 1 and Fig 1C), we found no further increase with EV from obese animals (Fig 2D). Collectively, these results show that the heightened effect of adipocyte EV observed in obesity does not depend on increased FAO protein transfer.

**Figure 2:**
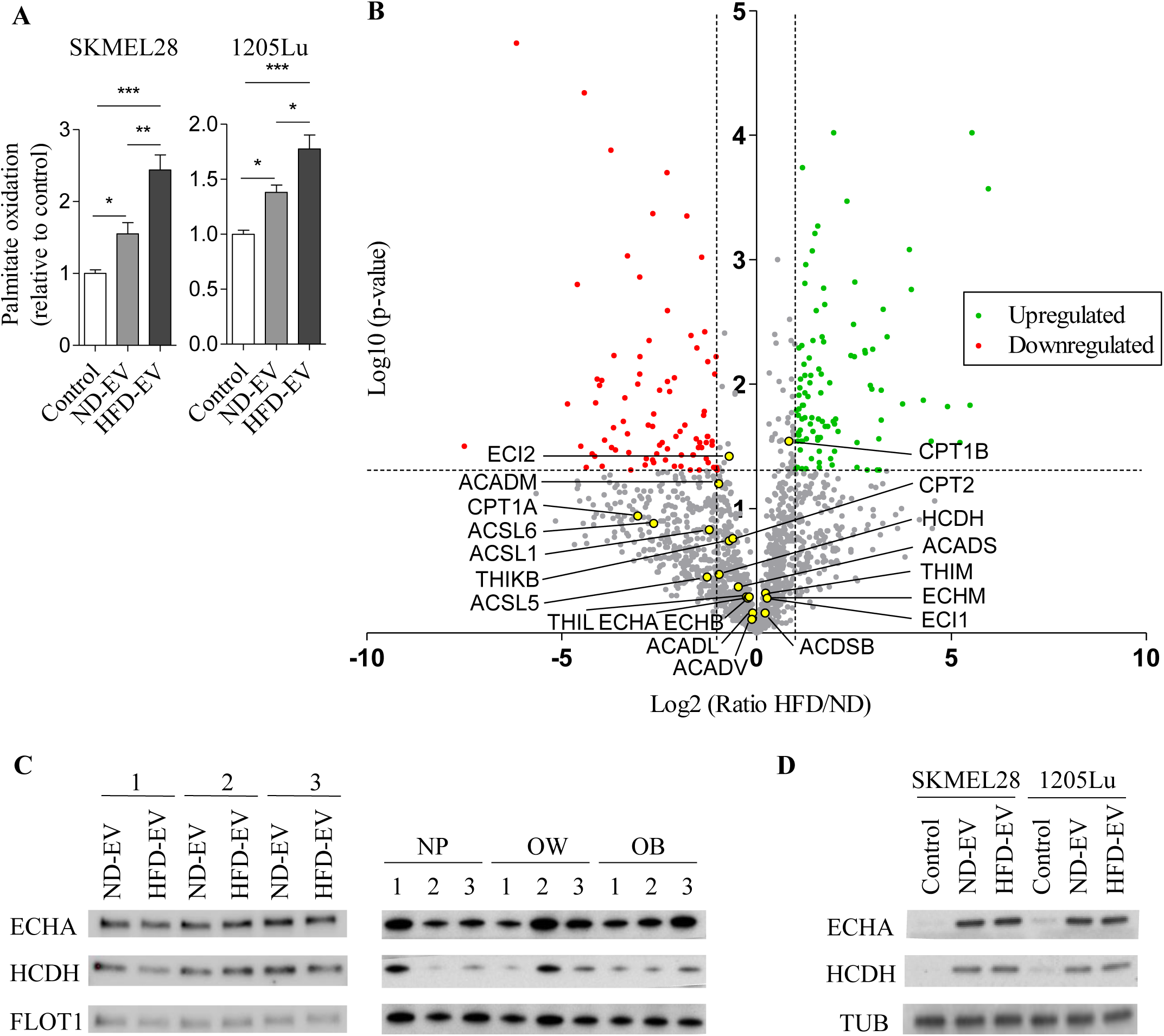
Adipocyte EV-induced FAO is increased by obesity, but this process is not dependent on increased protein transfer. A) SKMEL28 and 1205Lu cells were exposed, or not, to the indicated adipocyte EV from primary adipocytes from lean (ND) or obese (HFD) mice, then FAO was measured. Bars and error flags represent means ± SEM (n ≥ 5); statistically significant by one-way ANOVA with post hoc Tukey’s test, *P < 0.05, **P < 0.01, ***P < 0.001. B) Volcano plot of mass spectrometry-based quantitative proteomics results showing relative abundance of proteins in primary adipocyte EV from obese mice (HFD), as compared to those from lean mice (ND). The dashed lines indicate cut-off values and points colored in grey indicate proteins that display non-significant fold-change by Welch t-test between both conditions (n=3 for ND, 4 for HFD). Proteins involved in FAO are indicated by yellow dots. C) Western blot analysis of the indicated FAO enzymes in the EV secreted by primary adipocytes from lean (ND) and obese (HFD) mice (left panel) and from human individuals with varying BMI (normoponderal, NP; overweight, OW and obese, OB) (right panel). For each blot, extracts from 3 independent batches of murine samples or 3 independent individuals for human samples (1-3) are shown. Flotillin 1 (FLOT1) is used as a loading control. D) Western blot analysis of the indicated FAO enzymes in melanoma cells treated, or not, with EV from lean (ND) and obese (HFD) mice. Tubulin (TUB) is used as a loading control.

### Adipocyte EV transfer FA to melanoma cells to fuel FAO, a process increased by obesity

Increased FAO requires both the protein machinery necessary to perform the enzymatic processes and the presence of the substrate. So, we postulated that adipocyte EV transfer FA to melanoma cells and that this process may be amplified in obesity, which could account for increased tumor cell aggressiveness. EV secreted by mature 3T3-F442A adipocytes contain high levels of FA compared to their precursors (Fig 3A). To determine whether these FA are subsequently transferred to tumor cells, we used a lipid pulse-chase assay (Fig 3B). We loaded 3T3-F442A adipocytes with BODIPY FL C_16_ (Supplementary Fig 2A) and monitored the EV transfer of this fluorescent FA. We detected fluorescent FA in tumor cells exposed to adipocyte EV (Fig 3C and Supplementary Fig 2B). This fluorescent FA serves as a substrate for FAO in melanoma cells, as treatment with the FAO inhibitor, Etomoxir, leads to a further accumulation of fluorescent lipids (Fig 3D). We confirmed this finding using EV secreted by primary adipocytes (Supplementary Fig 2C). To determine whether FA transfer increases with EV from primary obese adipocytes, we evaluated FA content in EV secreted by primary murine and human adipocytes. FA content significantly increased in murine adipocyte EV in obesity (Fig 3E, top panel) and positively correlated with BMI (body mass index) in human samples (Fig 3E, bottom panel), although triglyceride content remained unchanged in obesity (Supplementary Fig 2D), consistent with recent lipidomic data (Flaherty et al, 2019). Since hypertrophic adipocytes internalize less lipids than their smaller counterparts (Frayn, 2001; Hill et al, 2009), (Supplementary Fig 2E), we could not use the pulse chase assay described in figure 3B to compare FA transfer by lean and obese adipocyte EV. Nevertheless, obese adipocyte EV strongly increased the total neutral lipid content in melanoma cells compared to lean adipocyte EV, which supports our hypothesis (Fig 3F). Thus, these results demonstrate that adipocytes transfer FA to melanoma cells through EV, through a process that is amplified by obesity.

**Figure 3:**
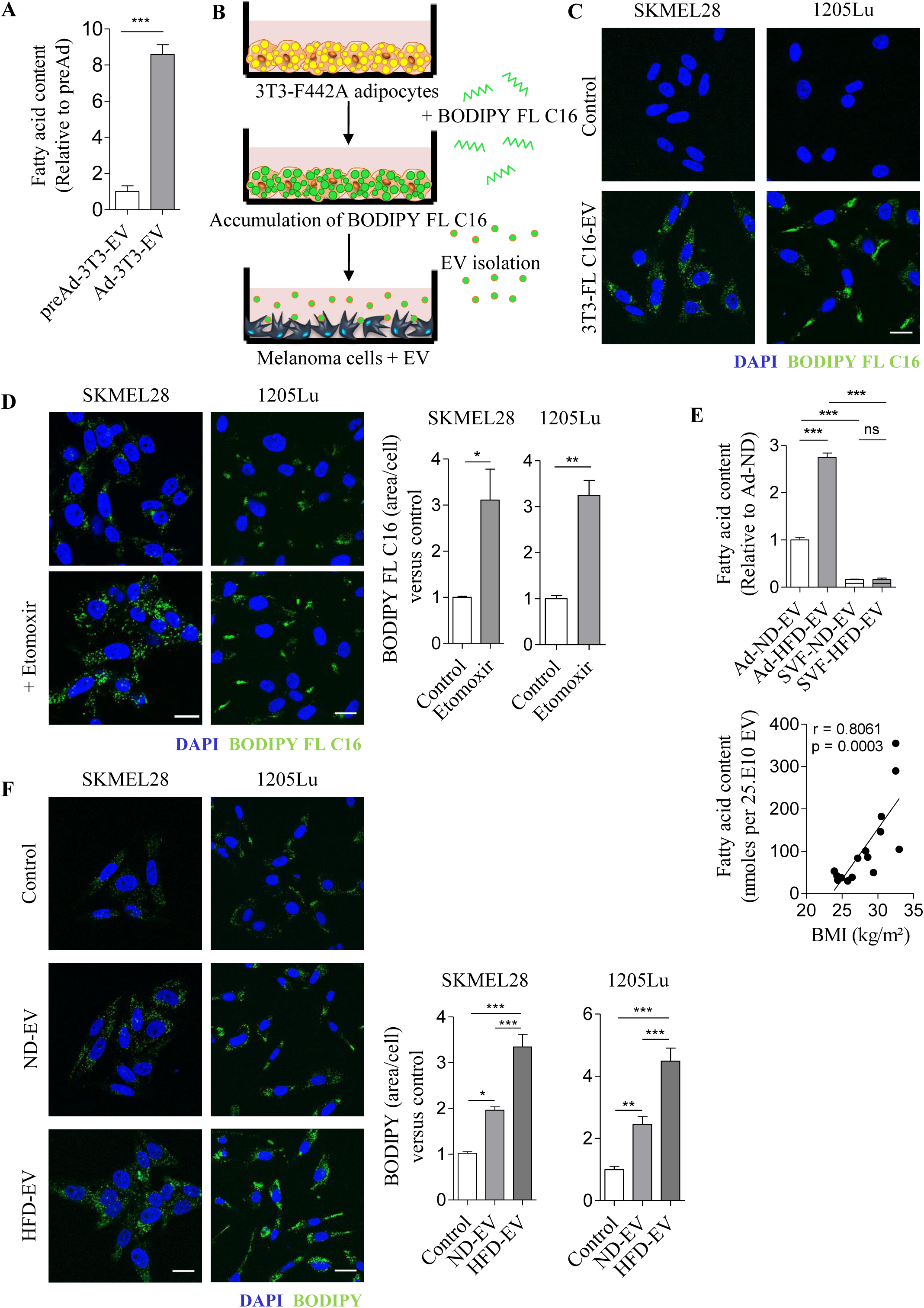
Adipocyte EV transfer FA to melanoma cells to fuel FAO, and this transfer is increased in obesity. A) Lipids were extracted from EV secreted by 3T3-F442A pre-adipocytes and differentiated 3T3-F442A adipocytes (respectively pre-Ad-3T3-EV and Ad-3T3-EV), and FA content was measured (n=6). B) Workflow of the assay used to evaluate FA transfer from adipocyte EV to melanoma cells. Mature 3T3-F442A adipocytes were loaded with BODIPY FL C16. Cells were then washed and fresh medium was added. 72h later, conditioned medium was harvested, and EV were isolated and added to melanoma cells. C) Indicated melanoma cells were incubated with EV from 3T3-F442A adipocytes previously loaded with BODIPY FL C16 (3T3-FL C16-EV) and, 24h later, cells were fixed and nuclei were counterstained with DAPI. D) Left panel, indicated cells were incubated with EV from 3T3-F442A adipocytes previously loaded with BODIPY FL C16 and immediately treated, or not, with Etomoxir for 24h. Then, cells were fixed and nuclei were counterstained with DAPI. Right panel, quantification of BODIPY FL C16 staining area per cell (n=3). E) Lipids were extracted from EV secreted by adipocytes from lean (ND) and obese (HFD) mice (n=5) (left panel) or from human adipose tissue samples from patients with varying BMI (right panel) and FA content was measured. F) Indicated cells were exposed, or not, to adipocyte EV from primary adipocytes from lean (ND) or obese (HFD) mice for 24h. Then, cells were fixed, stained with BODIPY and counterstain with DAPI. Left panel, Confocal microscopy observation. Right panel, quantification of BODIPY staining area per cell (n=5 for SKMEL28 and n=6 for 1205Lu). Scale bars represent 20µm. Bars and error flags represent means ± SEM; statistically significant by unpaired Student’s t-test (A, D), or by one-way ANOVA with post hoc Tukey’s test (E-F) whereas Spearman’s rank correlation was used to evaluate the correlation between FA content in human adipocyte EV and patient BMI (E). *P < 0.05, **P < 0.005. ***P < 0.001, ns: non-significant.

### Transferred FA are stored in lipid droplets and released by lipophagy

Although the FA transferred to melanoma cells by adipocyte EV fuel FAO, we also show an increase in neutral lipid storage within cytoplasmic lipid droplets (Fig 3F), a process known to prevent lipotoxicity (Listenberger et al, 2003). We therefore examined whether these stored lipids are mobilized to drive FAO. We observed that melanoma cells incubated with adipocyte EV present double membrane structures, characteristic of autophagosomes that contain lipids within their lumen (Fig 4A and Supplementary Fig 3A). Thus, we hypothesized that the transferred FA were degraded by lipophagy, an autophagic process that releases FA from lipid droplets (Singh et al, 2009). In accord with this theory, FA transferred by adipocyte EV colocalize with lysosomes (Fig 4B and Supplementary Fig 3B). We found that inhibiting autophagy using Bafilomycin A1 prevented the degradation of lipid stores accumulated in response to adipocyte EV (Supplementary Fig 3C) and blocked their pro-migratory effect on melanoma cells (Supplementary Fig 3D). We obtained similar results using the selective lysosomal acid lipase inhibitor, Lalistat 2 (Hamilton et al, 2012; Rosenbaum et al, 2010). This compound increased colocalization between fluorescent FA transferred by adipocyte EV and lysosomes (Fig 4B) and effectively inhibited degradation of transferred FA (Fig 4C and Supplementary Fig 4A). This result was also reflected in the total neutral lipid stores accumulated in the presence of adipocyte EV (Supplementary Fig 4B). We also found that lipophagy is requisite to elicit the effect of adipocyte EV on melanoma migration and motility (Fig 4D and Supplementary Fig 4C). Furthermore, inhibiting lipophagy also prevented degradation of stored lipids following exposure to both lean and obese primary adipocyte EV (Fig 4E and Supplementary Fig 4D), as well as their effect on cell motility (Fig 4F). Our results from the SILAC experiment revealed that various autophagic and lysosomal proteins are transferred by adipocyte EV to melanoma cells. This suggests that EV may promote lipophagy by providing these proteins (Supplementary Table 1). Thus, our results show that adipocyte EV-derived lipids stored by melanoma cells are released by lipophagy, which is crucial for the pro-migratory effect of EV.

**Figure 4:**
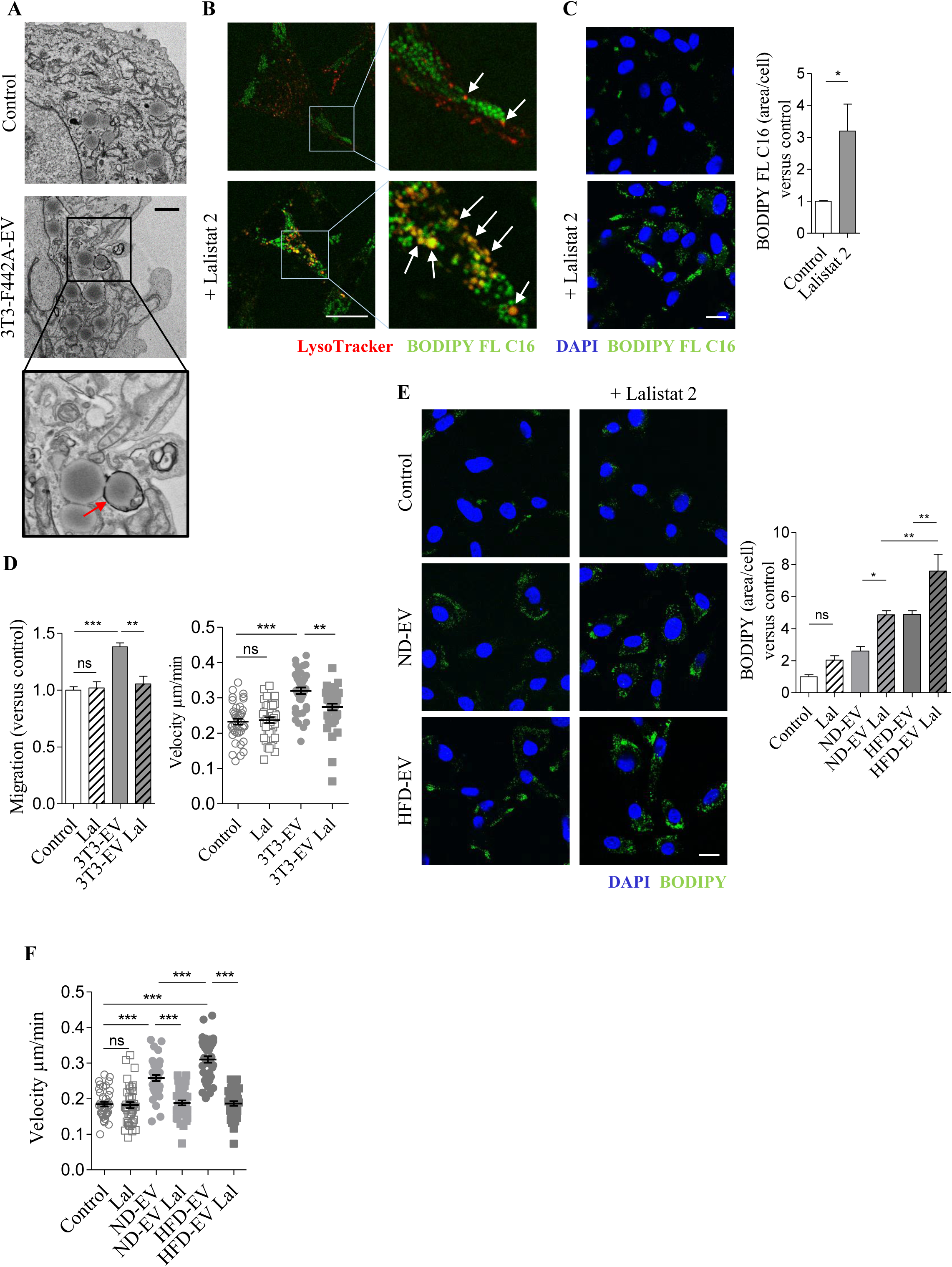
FA transferred from adipocytes to melanoma cells by EV are released from lipid droplets by lipophagy. A) Transmission electron micrographs of 1205Lu cells exposed, or not, to 3T3-F442A EV. A zoomed crop of the area with autophagic structures containing lipids (indicated by an arrow) is shown. Scale bar represents 1µm. B) 1205Lu cells were incubated with EV from 3T3-F442A adipocytes, previously loaded with BODIPY FL C16 in the presence, or not, of Lalistat 2. Then, live cells were stained with the LysoTracker probe and observed by confocal microscopy. C) 1205Lu were incubated with EV from 3T3-F442A adipocytes, previously loaded with BODIPY FL C16 and treated, or not, with Lalistat 2. Then, cells were fixed and counterstained with DAPI. Quantification of BOBIPY FL C16 staining per cell is shown beside (n=4). D) Left panel, 1205Lu cells were exposed to 3T3-F442A EV (3T3-EV) and treated, or not, with Lalistat 2 (Lal). Cell migration was then evaluated in Boyden chamber assays (n=5). Right panel, 3T3-F442A EV were added to 1205Lu cells, which were immediately treated, or not, with Lalistat 2. Cell motility was then tracked by video microscopy (n=40 cells per group). E) 1205Lu cells were exposed, or not, to adipocyte EV from lean (ND) or obese (HFD) mice with, or without, Lalistat 2 (Lal). Cells were then fixed, stained with BODIPY and counterstain with DAPI. Quantification of BODIPY staining per cell is shown on the right (n=5). F) 1205Lu cells were exposed, or not, to adipocyte EV from ND or HFD mice with, or without, Lalistat 2 (Lal). Cell motility was then tracked by video microscopy (n=40 cells per group). On all confocal microscopy images, scale bars represent 20µm. Bars and error flags represent means ± SEM; statistically significant by unpaired Student’s t-test (C), or by one-way ANOVA with post hoc Tukey’s test (D-F). *P < 0.05, **P < 0.005. ***P < 0.001, ns: non-significant.

### Adipocyte EV remodel the mitochondrial network to support melanoma migration

Increased lipid accumulation (Zhang et al, 2011) and subsequent increased FAO (Kuzmicic et al, 2014) can remodel the mitochondrial network. Interestingly, mitochondrial dynamics also play a crucial role in tumor cell migration (Altieri, 2017). Redistributing mitochondria toward cell extremities during tumor cell migration increases energy supply for actin polymerization in these zones (Cunniff et al, 2016). Consistent with such a process in response to adipocyte EV, treated melanoma cells adopt an elongated morphology and present profound changes in mitochondrial distribution switching from a perinuclear localization to a distribution throughout the cytoplasm, particularly toward membrane protrusions (Fig 5A). In our SILAC experiment, we found that key regulators of mitochondrial dynamics, FIS1 and OPA1, are among the transferred proteins from adipocytes to melanoma cells by EV (Supplementary Table 1). Moreover, increased melanoma cell migration and motility following treatment with adipocyte EV is completely abrogated by the mitochondrial fission inhibitor Mdivi-1 (Fig 5B-C and Supplementary Fig 5A). We confirmed these findings using gene-silencing experiments. Two different siRNAs targeting the key mitochondrial fission regulator DRP1 (DNM1L) effectively reduced the expression of the protein in melanoma cells (Fig 5D, left panel) and strongly abrogated the pro-migratory effect of adipocyte EV (Fig 5D, right panel). Finally, FAO was clearly required for mitochondrial redistribution in both lean and obese conditions, since mitochondria retain a perinuclear localization in the presence of Etomoxir (Supplementary Fig 5B).

**Figure 5:**
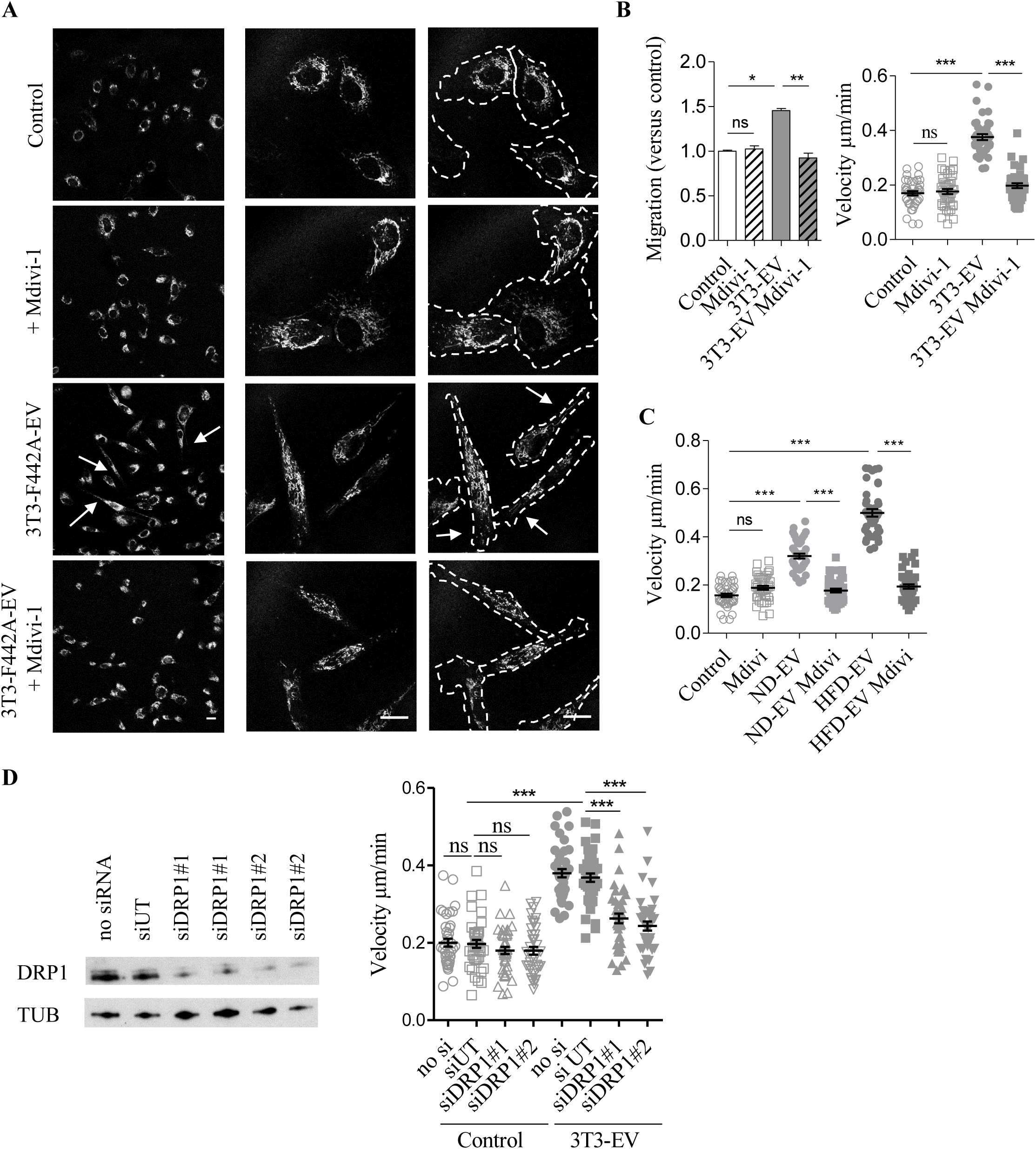
Adipocyte EV modify melanoma mitochondrial dynamics, a process that promotes melanoma aggressiveness and is exacerbated by obesity. A) 1205Lu cells exposed to 3T3-F442A EV and treated, or not, Mdivi-1, were stained with a MitoTracker probe, fixed and observed by confocal microscopy. Arrows indicate the presence of mitochondria in membrane protrusions. On the images to the right, the outline of cells is shown in dotted lines. Scale bars: 20µm. B) Left panel, 1205Lu cells were exposed to 3T3-F442A EV (3T3-EV) and treated, or not, with Mdivi-1. Cell migration was then evaluated in Boyden chamber assays (n=6). Right panel, 3T3-F442A EV were added to 1205Lu cells, which were immediately treated, or not, with Mdivi-1. Cell motility was then tracked by video microscopy. C) 1205Lu cells were exposed to adipocyte EV from lean (ND) or obese (HFD) mice and immediately treated, or not, with Mdivi-1. Cell motility was then tracked by video microscopy (n = 40 cells per group). D) 1205Lu cells were transfected, or not, with 2 different siRNA targeted against DRP1 (siDRP1 #1 or 2) or an untargeted siRNA (siUT). Left, 48h after transfection, protein extracts were prepared and DRP1 expression was evaluated by western blot. Right, 36h post transfection, cells were exposed to 3T3-F442A EV (3T3-EV) and cell motility was tracked by video microscopy (n = 40 cells per group). Bars and error flags represent means ± SEM; statistically significant by one-way ANOVA with post hoc Tukey’s test (B-D). *P < 0.05, **P < 0.005. ***P < 0.001, ns: non-significant.

In obesity, the amplified effect of adipocyte EV on cell motility also depends on mitochondrial fission, as Mdivi-1 totally abrogates their effect (Fig 5C). For FAO to efficiently take place in cell protrusions and supply energy for cell migration, we expected that both the mitochondrial machinery and the substrate would be present. In melanoma cells incubated with adipocyte EV, we found a striking lipid droplet redistribution toward cell protrusions (Fig 6A). Obesity further exacerbated this redistribution (Fig 6B). Moreover, they are found in close proximity with mitochondria in these areas, as observed with confocal microscopy (Fig 6C and Supplementary Fig 6A) and transmission electron microscopy (Fig 6D and Supplementary Fig 6B). Thus, these results show that mitochondrial dynamics, and possibly lipid droplet dynamics, trigger cell migration in response to adipocyte EV, with a heightened effect in obesity.

**Figure 6:**
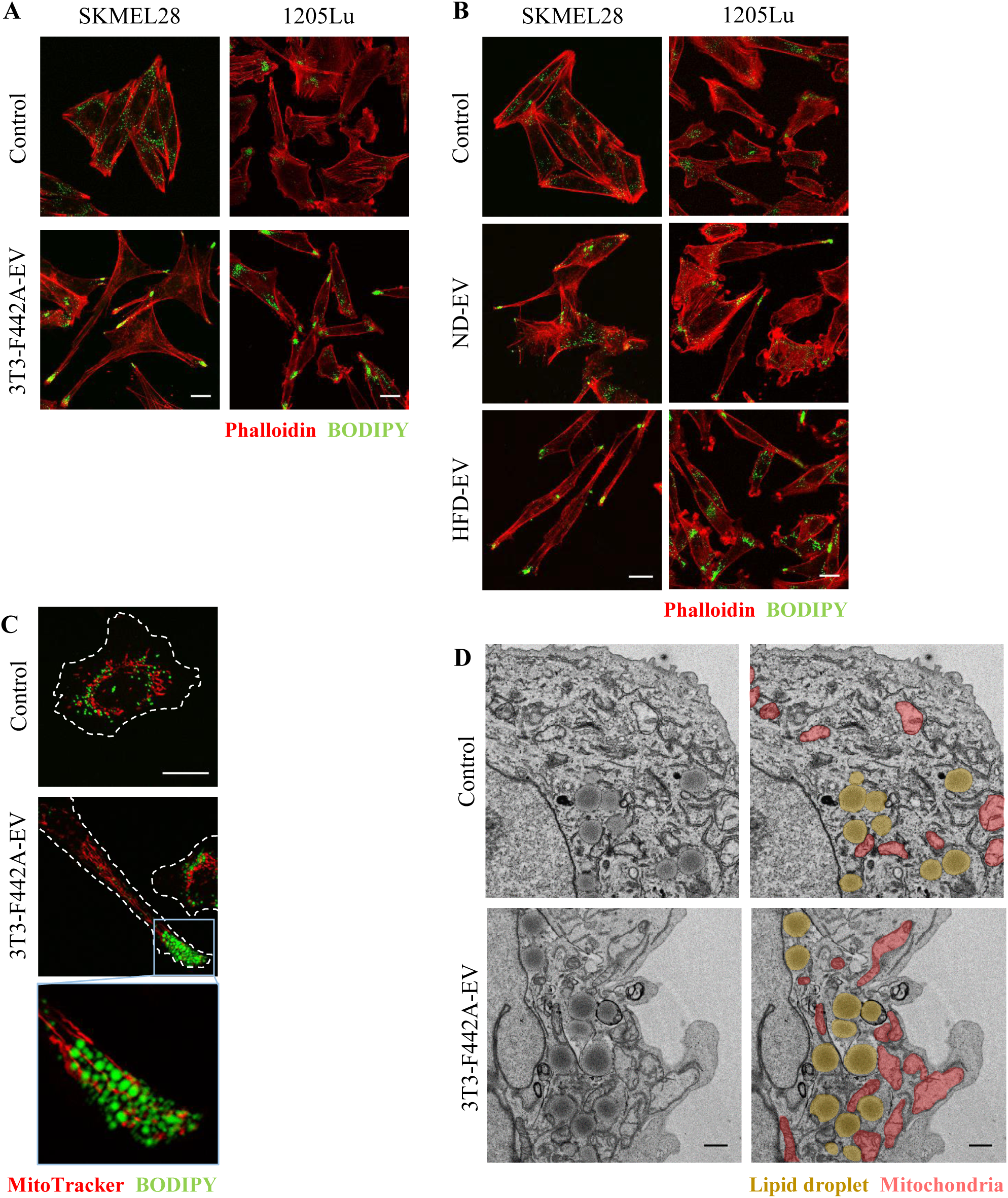
Lipid droplets are found in membrane protrusions, at proximity to mitochondria, in melanoma cells exposed to adipocyte EV. A) Melanoma cells exposed to 3T3-F442A EV were fixed, stained with BODIPY and Phalloidin. B) Melanoma cells exposed to EV secreted by adipocytes from lean (ND) or obese (HFD) mice were fixed, stained with BODIPY and Phalloidin. C) 1205Lu cells exposed to 3T3-F442A EV were stained with a MitoTracker probe. Then, cells were fixed and stained with BODIPY. A zoomed crop of the area containing mitochondria and lipid droplets in a membrane protrusion is shown. D) Transmission electron microscope observations of 1205Lu cells exposed, or not, to 3T3-F442A EV. Mitochondria are colored in red and lipid droplets are colored in yellow on images on the right. Scale bars represent 20µm for confocal microscopy images and 1µm for electron microscopy images.

### FAO and mitochondrial dynamics correlate with melanoma aggressiveness

To assess the clinical relevance of our findings, we performed a targeted analysis of human melanoma sample data that is publicly available online in The Cancer Genome Atlas database (TCGA, http://www.cbioportal.org). Although this tumor collection is not annotated with biometric data to study obesity, we investigated whether the cellular processes we identified here are key to melanoma progression in the general population. This analysis revealed that increased expression of FAO enzymes correlated with poor overall survival (OS) of melanoma patients (Fig 7A). Accordingly, neutral lipid content and FAO progressively increased in melanoma cell lines with increasing metastatic potential (Lazar et al, 2015), (Goodall et al, 2008) (Fig 7B-C). Finally, mitochondrial dynamics strongly correlated with melanoma progression, as heightened expression of genes involved in mitochondrial fission (FIS1) or fusion (MFN2 and OPA1) correlated with decreased OS in melanoma patients (Fig 7D).

**Figure 7:**
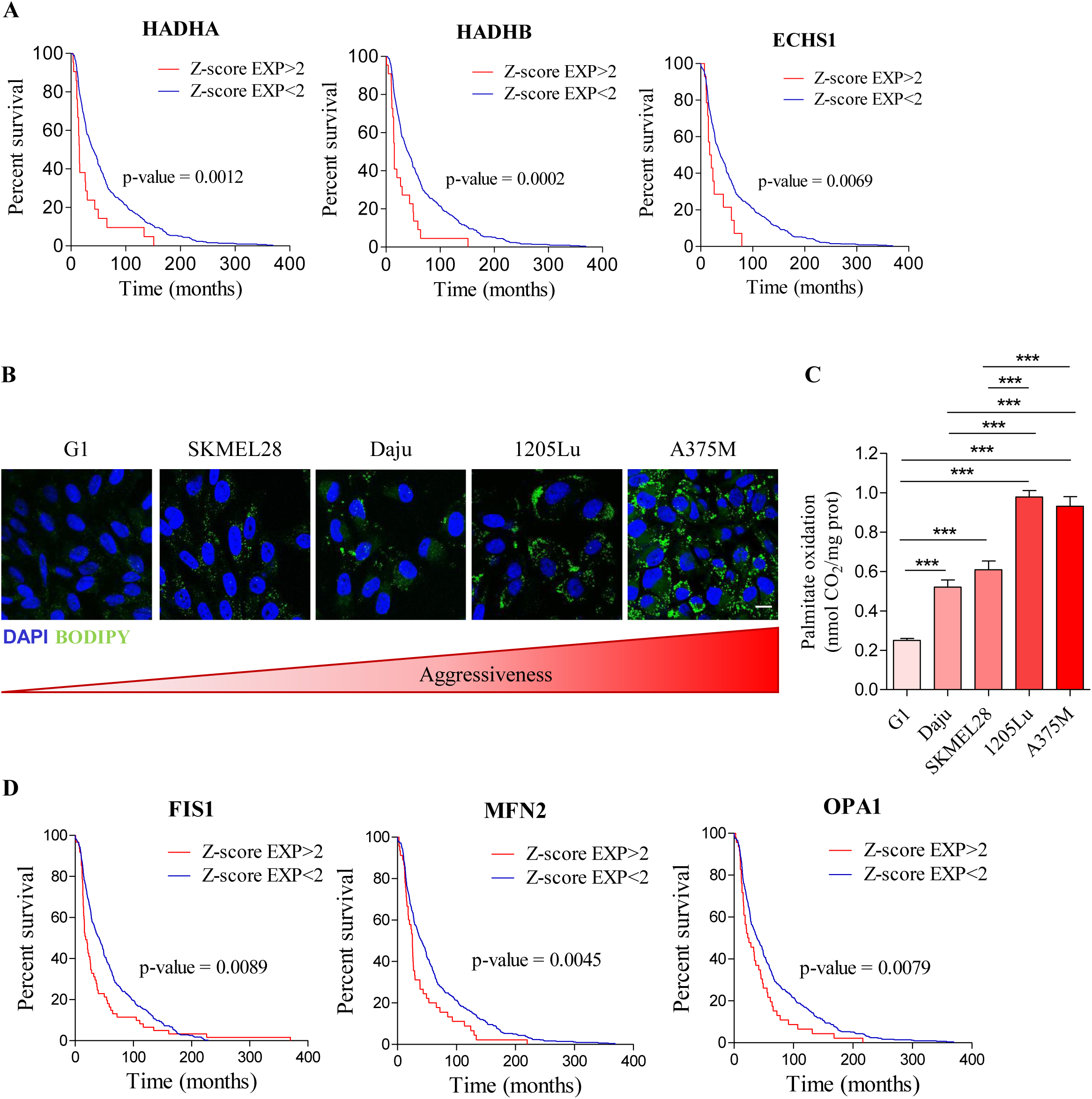
FA metabolism and mitochondrial dynamics are associated with melanoma aggressiveness. A) TCGA data were analyzed to reveal the effect of high mRNA levels of key FAO enzymes on the overall survival of patients with melanoma. Statistical significance of survival was evaluated with a Log-rank (Mantel-Cox) Test. B) BODIPY staining of neutral lipid stores in the indicated melanoma cell lines, which present increasing degrees of aggressiveness (from left to right). DAPI was used to counterstain nuclei. Scale bar: 20µm. C) FAO in the indicated melanoma cell lines was measured. Bars and error flags represent means ± SEM (n=5); statistically significant by one-way ANOVA with post hoc Tukey’s test, ***P < 0.001. If not indicated, non-significant. D) TCGA data were analyzed to reveal the effect of high mRNA levels of key actors in mitochondrial dynamics on the overall survival of patients with melanoma. Statistical significance of survival was evaluated with a Log-rank (Mantel-Cox) Test.

## Discussion

In summary, we reveal the importance of EV in the metabolic dialog between adipocytes and melanoma cells, which promotes FAO and, ultimately, tumor aggressiveness. Previous studies showed such a process occurred at the invasive front of tumors where secretions from cancer cells provoke adipocyte lipolysis, which provides tumors with the necessary substrate for FAO. However, our findings demonstrate that EV secreted by naïve adipocytes provoke a similar process and, in fact, provide melanoma cells with both the required machinery (proteins) and the substrate (FA) for FAO. We determined the proteins transferred from adipocytes to tumor cells by EV using a groundbreaking approach based on the SILAC technique, which permitted widespread labeling of adipocyte proteins for subsequent identification within tumor cells by mass spectrometry. To our knowledge, this is the first time that a comprehensive analysis of all of the proteins transferred between two cell types by EV has been performed. We also illustrate the utility of this technique for any cultured cell types to study EV-mediated intercellular protein transport. Using this approach, we found that only approximately 30% of labeled adipocyte EV proteins are detected in recipient melanoma cells, with some highly abundant EV proteins excluded from transfer. This process suggests a highly selective EV-mediated protein transfer between adipocytes and melanoma cells. This could be the result of some populations of EV not being internalized by melanoma cells or some EV or the proteins they carry being rapidly sorted for degradation after internalization. While we do not fully understand how EV are selectively internalized and/or processed in recipient cells, our approach provides a foundation to address these questions in future studies. Despite this selective transfer, melanoma cells internalize adipocyte proteins involved in FAO, mitochondrial respiration and ATP production through EV. Importantly, we also identified key contributors in other cellular processes induced in melanoma cells in response to adipocyte EV that help regulate their pro-migratory effects, lipophagy and mitochondrial dynamics (Supplementary Table 1).

In addition to this protein transfer, we demonstrated that adipocyte EV also convey FA to tumor cells to drive FAO. Increases in this process heightened the effect of adipocyte EV in obesity. Shuttling of metabolic substrates from the tumor microenvironment to cancer cells by EV has previously been described. Indeed, CAF-derived EV transport metabolites, such as amino acids, lipids and TCA cycle intermediates, to tumor cells, which serves as triggers for central carbon metabolism (Zhao et al, 2016). Even though we know FA are released by adipocytes in response to tumor secretions for transfer to melanoma cells (Kwan et al, 2014; Zhang et al, 2018), we reveal that naïve adipocytes, that have never encountered tumor secretions, can also convey lipids to cancer cells through EV. Thus, adipocyte EV mediate a metabolic cooperation between adipocytes and melanoma cells, acting as shuttles to convey both the protein machinery and the lipid substrate required for FAO.

Analysis of TCGA data revealed that high levels of FAO enzymes correlate with lower OS in melanoma patients (Fig 7A). Consistent with this, studies found upregulated genes associated with FAO in aggressive melanoma (Rodrigues et al, 2016; Xu et al, 2012). Our work demonstrates that FAO depends on both the endogenous traits of melanoma cells and a metabolic cooperation with adipocytes mediated by EV. Although we did not observe an increased transfer of proteins involved in FAO in obese conditions, we cannot exclude the relevance of these enzymes under physiological conditions. Obese adipocytes do secrete larger quantities of EV compared to lean counterparts, which may further increase their effect in cancer patients (Flaherty et al, 2019; Lazar et al, 2016). In experimental conditions using equal quantities of vesicles, obese adipocyte EV convey higher levels of FA, which increases lipid accumulation in melanoma cells. In obesity, although stimulated lipolysis induced by activation of adrenergic receptors decreases, basal lipolysis is heightened (Duncan et al, 2007; Verboven et al, 2018). This process may increase FA release and, consequently, a higher availability of these substrates for tumor cells. Based on our results, we conclude these FA are partially secreted through EV.

Understanding the mechanisms that promote tumor progression in obese patients is crucial to develop relevant therapies. Here, we show that a metabolic cooperation exists between adipocytes and tumor cells and is amplified in obesity, suggesting that obese cancer patients would likely benefit from FAO inhibitors and/or molecules that inhibit EV uptake. As EV diffuse through tissues and circulate throughout the organism, they may influence tumors not only at proximity to adipose depots, but also at distance. Adipocyte EV may also accompany circulating tumor cells to provide them with the nutrients they require to reach and colonize metastatic niches. As this study focuses on naïve adipocyte EV, our results suggest these vesicles likely play significant roles in metabolic processes in a physiological context. Elegant studies demonstrated the importance of EV in the dialog between adipocytes and endothelial cells (Crewe et al, 2018), as well as AT macrophages (Flaherty et al, 2019). In light of our results, we also postulate that adipocyte EV can regulate energy metabolism, for example by providing muscles with the necessary molecules for FAO during physical activity.

We also identified the cellular processes induced in melanoma cells that process lipids transferred from adipocyte EV that then trigger metabolic remodeling and cell migration. We found that FA transferred by adipocyte EV are stored in lipid droplets and are then mobilized by melanoma cells through lipophagy. Lipophagy inhibitors prevented the degradation of stored lipids in response to adipocyte EV in both lean and obese conditions. Although the involvement of lipophagy in melanoma metabolism remains unclear, a recent study found that melanospheres containing cancer stem cells store lipids mobilized during differentiation of these cells by lipophagy (Giampietri et al, 2017).

Transferred FA that fuel FAO may remodel the mitochondrial network in melanoma cells to redistribute these organelles to the cell extremities, which promotes cell migration. The importance of mitochondrial dynamics in tumor cell migration is increasingly recognized (Altieri, 2017). Although mitochondrial fission is a focus of many studies, both fission and fusion events are critical to redistribute mitochondria towards the leading cell edge during migration (Desai et al, 2013). However, little is known about mitochondrial dynamics in melanoma progression. The limited data available suggests that the mitochondrial network is reorganized during B16-F10 migration (Cunniff et al, 2016). Consistent with this, our analysis of TCGA data revealed that increased mRNA expression of key actors involved in mitochondrial dynamics correlates with lower OS in melanoma patients. Inhibiting mitochondrial fission abrogates the increase in cell motility observed after treatment of melanoma cells with adipocyte EV, which highlights the importance of this process. Importantly, we linked mitochondrial redistribution to EV-induced FAO, as Etomoxir prevented mitochondrial relocalization to membrane protrusions. We identified key regulators of mitochondrial dynamics among the proteins transferred from adipocytes to melanoma cells by EV. Although the changes in the mitochondrial network that we found in the presence of adipocyte EV require FAO, we acknowledge other substrates may facilitate this process. Interestingly, we also showed that lipid droplets also relocalize toward membrane protrusions in melanoma cells exposed to adipocyte EV. This suggests that substrate is brought directly to specific sites to continue to fuel FAO during melanoma migration. Such a proximity can occur in other models, in which it increases lipid droplet and mitochondria interactions to facilitate lipid transfer between the two organelles (Herms et al, 2015; Rambold et al, 2015).

In summary, our findings demonstrate that adipocytes EV provide both the machinery (enzymes) and substrate (FA) required for FAO to melanoma cells. The FA transferred by these EV are stored in lipid droplets and mobilized by lipophagy to fuel FAO. We identified mitochondrial dynamics that link adipocyte-induced FAO and tumor cell migration. We observed a mitochondrial redistribution toward cell protrusions that requires mitochondrial fission, which increases cell migration induced by adipocyte EV. In obesity, a higher supply of FA through EV amplifies this entire process (Fig 8). These results demonstrate a metabolic cooperation occurs between adipocytes and tumor cells. Further, we reveal the underlying cellular processes in this interaction. We propose these pathways could comprise novel therapeutic targets to treat obese melanoma patients.

**Figure 8:**
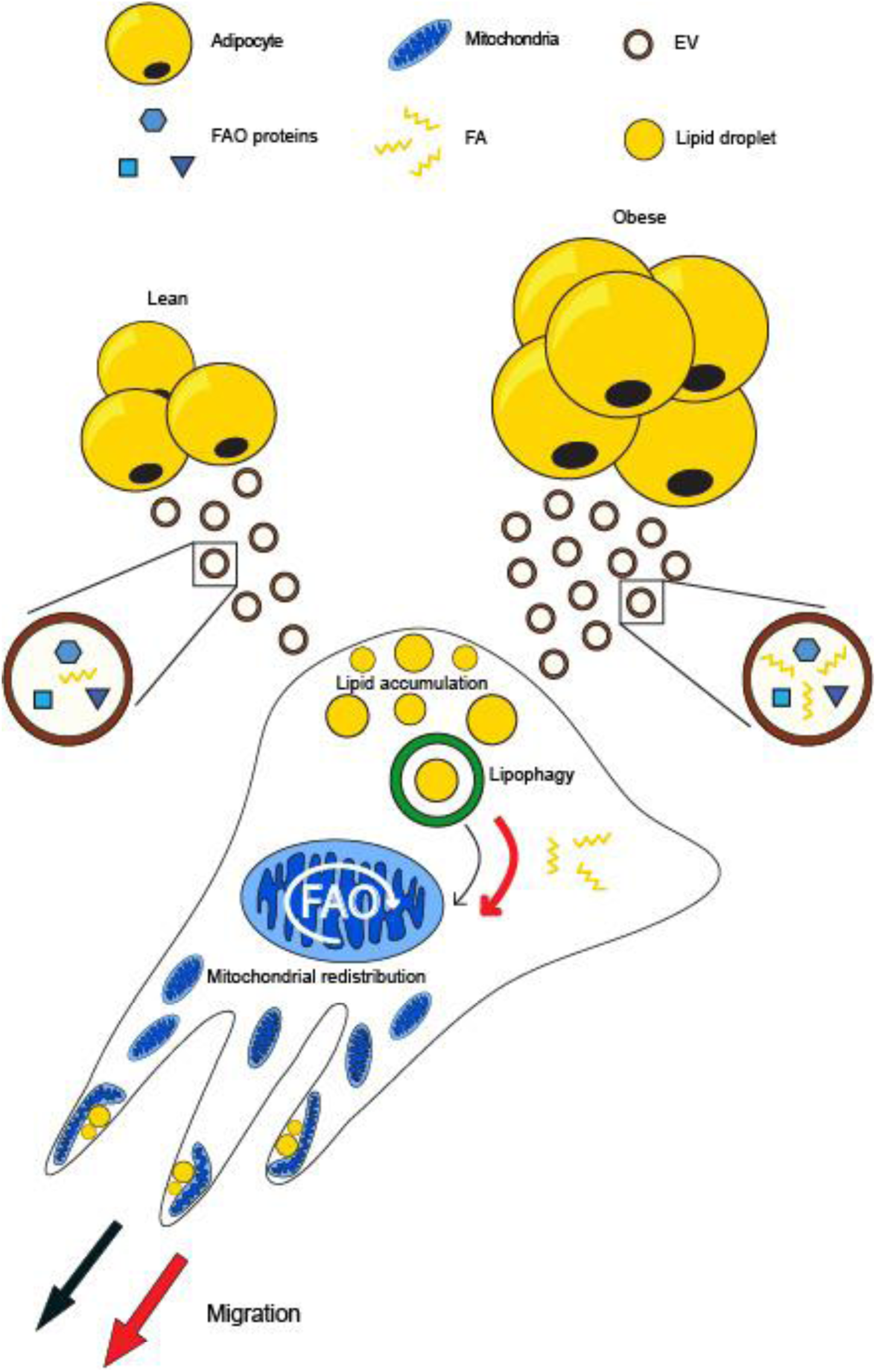
Molecular actors and cellular processes involved in the increased effect of adipocyte EV in obesity. Adipocytes release EV that transfer the machinery (enzymes) and the substrate (FA) required for FAO to melanoma cells. This leads to accumulation of lipid stores within tumor cells, which are degraded by lipophagy to release FA and fuel FAO. The increase in mitochondrial activity leads to a reorganization of the mitochondrial network. Indeed, in cells treated with adipocyte EV, these organelles, as well as lipid droplets, are redistributed towards tumor cell membrane protrusions. This process supports cell migration by providing the energy necessary for cytoskeletal remodeling that takes place in these areas during cell migration. In obesity, an increased supply of FA within adipocyte EV amplifies this entire process, ultimately further increasing tumor cell aggressiveness.

## Materials and methods

### Reagents and antibodies

BODIPY® 493/503 (hereafter referred to simply as BODIPY), BODIPY™ FL C16, MitoTracker Red CMXRos, LysoTracker Deep Red, DAPI and Rhodamine Phalloidin were obtained from Molecular Probes-Thermo Fisher Scientific (Eugene, OR, USA). [1-14C] palmitate was obtained from PerkinElmer (Waltham, MA, USA). Etomoxir, Mdivi-1, bovine serum albumin (BSA) and insulin were obtained from Sigma Aldrich (Saint Louis, MO, USA). Lalistat 2 was obtained from Tocris Cookson (Bristol, UK). Bafilomycin A1 was purchased from Euromedex (Strasbourg, France). Antibodies were obtained from the following sources: FLOT1 from Santa Cruz Biotechnology (Dallas, TE, USA) used at 1/200); ECHA and HCDH from Invitrogen-Thermo Fisher Scientific (Auckland, NZ) used at 1/1000; α-Tubulin (clone Ab-2) from Thermo Fisher Scientific (Eugene, OR, USA) used at 1/1000; DNM1L (DRP1), used at 1/500 from Sigma Aldrich. DharmaFECT1 transfection reagent and siRNA were obtained from Dharmacon™ (Cambridge, UK). siRNA sequences are as follows: Untargeted (UT) siRNA: 5’ UAGCGACUAAACACAUCAA-3’; siRNA DRP1#1: 5’ GGAGCCAGCUAGAUAUUAA; siRNA DRP1#2: 5’ CAUCAGAGAUUGUUUACCA.

### Cell lines, culture and treatments

SKMEL28 cells are an established line that originate from a human metastatic lymphonodal lesion (Carey et al, 1976). 1205Lu were derived from the vertical growth phase primary lesion-derived cell line WM793 by serial passage through athymic mice and selection of cells from lung metastases (Kath et al, 1991). Both melanoma lines were provided by Dr Lionel Larue (Institut Curie, Orsay, France). The murine 3T3-F442A preadipocyte cell line was purchased from the European Collection of Cell Cultures.

All cell lines were cultured in DMEM supplemented with 10% FCS, 125mg/mL streptomycin, and 125UI/mL penicillin (all purchased from GIBCO-Thermo Fisher Scientific, Eugene, OR, USA) and maintained at 37°C in a humidified atmosphere with 5% CO2. Cells were used within 2 months after resuscitation of frozen aliquots and regularly tested for mycoplasma contamination.

To obtain mature adipocyte cell line, 9.10^4^ 3T3-F442A preadipocytes were seeded in 6-well plates and, 3 days later, once cells had reached confluence, differentiation was induced by supplementing medium with 50nM insulin for 14 days. The term “adipocyte” refers to cells that were differentiated for at least 14 days. To condition medium, 1.5mL of ultracentrifugated DMEM (protocol for preparation below) were added per well for 72h.

For treatment of melanoma cells with EV in 6-well plates, for example, 1.10^5^ melanoma cells were seeded and, 24h later, 5.10^10^ 3T3-F442A or primary adipocyte EV were added per well. Cells were incubated with EV for 24h or 48h for experiments to study lipid accumulation, 48h for experiments to study cell metabolism and mitochondrial dynamics and 72h for experiments to evaluate cell migration. Where indicated, Etomoxir (50µM), Bafilomycin A1 (5nM) or Mdivi-1 (20µM) were added 24h before the end of incubation with EV, unless specified otherwise. Lalistat 2 (50µM) was added from the beginning of EV treatment.

For siRNA treatment, cells were transfected with 50nM siRNA in presence of DharmaFECT1, as indicated by the supplier. 6h post-transfection, fresh media and, if needed, EV, were added. Cells were used 48h post-transfection for western blot analysis and 36h post-transfection to track cell motility.

### Animals and primary cell isolatio

Mice were handled in accordance with National Institute of Medical Research (INSERM) principles and guidelines. All experiments were approved by the local committee on ethics of animal experimentation. Mice were housed according to national and institutional guidelines for human animal, in a controlled environment with a 12:12-h light-dark cycle. Eight week old C57BL/6J male mice (Janvier, Le Genest St Isle, France) were fed a normal or high fat diet (ND and HFD) for 15 weeks [ND (PicoLab Rodent Diet 20, Purina Mills Inc., Brentwood, MO, USA) and HFD (Research Diets Inc., New Brunswick, New Jersey, USA)]. The energy contents of the diets were as follows: 16% protein, 80% carbohydrate, and 4% fat for the ND; and 20% protein, 20% carbohydrate, and 60% fat for the HFD. After 15 weeks of diet, the average weight of ND mice was 29.6g (+/− 1.2g) and of HFD mice was 48.9g (+/−3.8g). Mice were then sacrificed to collect inguinal subcutaneous and visceral AT. AT was weighed and placed in DMEM supplemented with 1% BSA (2.5mL/g of tissue). Adipocytes and the stromal vascular fraction (SVF) were isolated as previously described (Lazar et al, 2016). To condition medium, one million adipocytes per mL were suspended in ultracentrifugated DMEM overnight (protocol for preparation below).

### Preparation of human adipocytes

Human AT samples were collected from abdominal dermolipectomies in accordance with the recommendations of the ethics committee of the Rangueil Hospital (Toulouse, France). All patients gave their informed consent in accordance with the Declaration of Helsinki Principles as revised in 2000. All of them were drug free and healthy (lean patients: 20<BMI (body mass indexes) <25; overweight patients 25<BMI<30; obese patients BMI>30). Tissues were processed within 1 hour of surgical resection. Briefly, AT was separated from skin and large blood vessels, glandular tissue and fascia were discarded. AT was then weighed, cut into <1mm^3^ pieces and adipocytes were isolated as previously described (Lazar et al, 2016). To condition medium, one million adipocytes per mL were suspended in ultracentrifugated DMEM overnight (protocol for preparation below).

### Preparation of ultracentrifugated DMEM, EV isolation and Nanoparticle Tracking Analysis (NTA)

To prepare medium to be conditioned for EV isolation, DMEM supplemented with 10% FCS was depleted of contaminating vesicles by overnight centrifugation at 100,000g. The conditioned medium was obtained from 3T3-F442A or primary adipocytes as described above, and small EV, enriched in exosomes, were purified as previously described (Lazar et al, 2015). We previously referred to the population of EV that we study as exosomes (Lazar et al, 2016), since we use a high-speed spin to isolate small EV, which are enriched in this type of vesicle (Thery et al, 2006). However, due to new guidelines from the International Society for Extracellular Vesicles (Gould & Raposo, 2013; Lotvall et al, 2014; van der Pol et al, 2016), we will refer to this same population simply as ‘EV’ in the present study. EV number and size distribution were analyzed by NTA with a NanoSight LM10-HS (Malvern, Orsay, France), equipped with a 405nm laser as previously described (Lazar et al, 2015).

### Stable isotope labeling with amino acids in cell culture (SILAC)

3T3-F442A preadipocytes were grown and differentiated in a modified DMEM designed for isotope labeling in cell culture [Sigma Aldrich (Saint Louis, MO, USA)] supplemented with 10% dialyzed FCS (GIBCO-Thermo Fisher Scientific, Eugene, OR, USA), 125mg/mL streptomycin, 125UI/mL penicillin, 50nM of insulin and 3.5g/L of glucose. The following amino acids were also added to the culture medium (200mg/L of each): L-lysine:2HCl (^13^C_6_), L-arginine:HCl (^13^C_6_) (both purchased from Cambridge Isotope laboratories Inc, Tewksbury, MA, USA), L-proline and L-leucine (Sigma). At maturation, adipocytes were incubated in ultracentrifugated DMEM for 72h before collecting conditioned medium for EV isolation. SKMEL28 cells were incubated with these EV for 12h. Labeled EV and cells treated or not with these EV were lysed in PBS containing 1% SDS and protein concentration was determined using the DC™ Protein Assay from Biorad. Samples were then processed and analyzed using nano-LC-MS/MS, as described in supplementary material and methods.

### Comparative proteomic analysis of EV from lean and obese adipocytes

EV released by adipocytes from lean or obese mice were isolated and purified as described earlier. EV were lysed in PBS containing 1% SDS and protein concentration was determined using the DC™ Protein Assay from Biorad. Samples were then processed and analyzed using nano-LC-MS/MS, as described in supplementary material and methods. Three ND and four HFD independent EV preparations were analyzed with technical duplicates for each.

### BODIPY, Rhodamine Phalloidin and DAPI staining in fixed cells

Cells were seeded on glass coverslips and, after the indicated treatments, were fixed with 3.7% paraformaldehyde for 15min at room temperature and permeabilized with 0.2% Triton X-100 for 5min. Then, cells were incubated with BODIPY (used at 1μg/ml) for 15min, the rhodamine phalloidin probe (used at 6.6µM) for 1h and/or DAPI (used at 1μM) for 5min. Coverslips were mounted using VECTASHIELD® Antifade Mounting Medium (Vector Laboratories, Burlingame, CA, USA).

### BODIPY, MitoTracker and LysoTracker staining in live cells

Cells were seeded in Lab-Tek® chamber slides, or glass coverslips for analysis of mitochondrial redistribution, and treated as indicated. Cells were then washed twice with prewarmed PBS (with 1%BSA for BODIPY and LysoTracker staining) and incubated with BODIPY (1µg/ml in culture medium with no FCS, 15min), MitoTracker Red CMXRos (75nM in PBS, for 45min) or LysoTracker Deep Red (50nM in PBS with 1% BSA, for 1h) at 37°C. Cells were then washed three times with pre-warmed PBS (with 1% BSA after BODIPY and LysoTracker staining) then fixed (for analysis of mitochondrial redistribution) or incubated in pre-warmed medium with appropriate treatments for analyses in live cells.

### FA extraction and dosage

For lipid extraction, 25.10^12^ EV were resuspended in 200µL of PBS, then 1.5mL of methanol was added and samples were vortexed. Five mL of Tert-Butyl methyl ether (MTBE) was added before vortexing again and then samples were incubated for 1h at room temperature under mixing. Finally, 1.2mL of water were added to separate aqueous and organic phases. The organic phase was evaporated under liquid nitrogen and lipids were resuspended in 50µL of fatty acid assay buffer (Free Fatty Acid Assay Kit, Abcam, Cambridge, MA, USA). FA (contained in 5µL of buffer) were dosed using this kit, as indicated by the supplier.

### FA transfer by 3T3-F442A adipocyte EV

3T3-F442A mature adipocytes were washed twice with prewarmed PBS with 0.1% BSA, then incubated with BODIPY FL C16 at 5µM in DMEM with no FCS and with 0.1% BSA and 50nM insulin for 6h. Adipocytes were washed twice with pre-warmed PBS with 0.1% BSA then incubated in ultracentrifugated DMEM with 50nM insulin for 72h before collecting conditioned medium. EV were isolated from the medium as described above, and then incubated with unstained melanoma cells.

### Measurement of FAO

Cells, treated or not with adipocyte EV, were incubated for 3h with 1mL of warmed (37°C), modified Krebs Ringer buffer pH 7.4 containing 1.5% FA-free BSA, 5mM glucose, 1mM palmitate (Sigma Aldrich), and 0.5μCi/ml ^14^C-palmitate (Perkin Elmer, Waltham, MA, USA) as previously described (Wang et al, 2017). Following this treatment, 1mL of incubation media was transferred to a glass tube and a microtube containing 300μL of NaOH 1N was placed in the vial to capture the ^14^CO_2_. The buffer was acidified with 1mL of 1M H_2_SO_4_, and the vial was sealed and left overnight before counting the radioactivity (CytoScint, MP Biomedicals, Illkirch Graffenstaden, France). All assays were performed in triplicates. Cells were lysed and protein concentration was determined using the DC™ Protein Assay from Biorad (Hercules, Ca, USA) for data normalization.

### Western blotting analysis

Cells or EV were lysed in PBS containing 1% SDS and protein concentration was determined using the DC™ Protein Assay from Biorad. 2μg of proteins were electrophoresed on SDS-PAGE. WB analysis were performed as previously described (Lazar et al, 2015).

### Confocal microscopy

A confocal microscope FV-1000 was used to observe cells with a 60X oil PLAPON OSC objective (Olympus, Hamburg, Germany). Images were processed to filter the noise with ImageJ software (National Institute of Health, Bethesda, MD) and a similar filter was used to analyze all acquisitions for the experiment. For the quantification of lipid accumulation, lipid droplet area per cell was quantified with ImageJ software. For each independent experiment, at least 4 images were analyzed to obtain a mean value. Graphs represent pooled means from at least 3 independent experiments.

### Migration assays

Equal numbers of cells (20.10^4^) in serum-free DMEM were added to the upper compartments of Transwell chambers (ThinCerts®, 12 wells, 8µm pores, Greiner Bio-One). DMEM supplemented with 10% FCS was added to the bottom chamber as a chemoattractant. Cells were then incubated at 37°C for 12h and 24h for 1205Lu and SKMEL28 cells respectively. Migrating cells were evaluated as previously described (Lazar et al, 2015).

### Video microscopy and analysi

Images of cells were acquired every 30min over 24h with an automated Leica DMIRB microscope equipped with a CoolSnap Ez camera (Roper Scientific, Lisses, France) with a 10X objective, programmed using MetaMorph software (Molecular Devices, San José, CA, USA). All images were processed using the same filter to reduce noise with ImageJ software. Then, cells were tracked in time-lapse image stacks using the Manual Tracking plugin in ImageJ to determine velocity.

### Transmission electron microscopy

Specimens were prepared as previously described (Lazar et al, 2015). Grids were examined with a transmission electron microscope (Jeol JEM-1400, Akishima, Tokyo, Japan) at 80kV.

### TCGA (The Cancer Genome Atlas) analysis

Analysis for the correlations between overall survival (OS) and mRNA expression of indicated genes was performed using the cBioPortal for Cancer Genomics Web site, http://www.cbioportal.org/public-portal/. The data set ‘Skin Cutaneous Melanoma (TCGA, Provisional)’, containing a total of 479 samples, was used to analyze the RNA Seq V2 RSEM z-score for each gene, which represents a normalized relative expression level. The z-score cut-off used was a 2-fold increase in mRNA expression. Data was downloaded and association of OS with increased gene expression was represented using Kaplan-Meier survival analysis and significance was assessed by Log-rank (Mantel-Cox) Test with Prism 5 software. Data were accessed from the cBioPortal in September 2018.

### Statistical analysis

Values are means ± standard error of the mean. The statistical significance of differences between the means (of at least three independent assays) was evaluated using Student’s t-tests, if 2 groups are compared, or one-way ANOVA, if more than 2 groups are compared, with the indicated associated post hoc tests using GraphPad Prism software. To determine the appropriate post hoc test to apply, normality of samples was determined using a Kolmogorov–Smirnov test. p values below 0.05 (*), <0.01 (**), <0.001 (**) and <0.0001 (***) were deemed as significant, ns: non-significant. Spearman’s rank correlation was used to evaluate the correlation between FA content in EV and BMI.

## Acknowledgements

The authors thank Lionel Larue for donating melanoma lines and Pascale Belenguer for insights and guidance on experiments relating to mitochondrial dynamics. Studies were supported by the ‘Ligue Régionale Midi-Pyrénées contre le Cancer’, ‘Fondation ARC pour la recherche sur le cancer’, and ‘Société Française de Dermatologie’ to LN and CM. The work was also supported by the Région Midi-Pyrénées, European Found (FEDER), and ‘Toulouse Métropole’, and the French Ministry of Research with the program “Investment for the Future, National Infrastructures for Biology and Health” (ProFI, Proteomics French Infrastructure project, ANR 10-INBS-08) to OBS. We thank ‘Région Midi-Pyrénées’ for supporting the ‘Toulouse Réseaux Imagerie’ platform. This work benefited of the assistance of the Multiscale Electron Imaging platform (METi) of the FRBT (Federation de Recherche en Biologie de Toulouse. EC is a recipient of a PhD fellowship from ‘Ligue nationale contre le Cancer’, and EC and IL are recipients of PhD fellowships from ‘Fondation ARC pour la recherche sur le cancer’. CA benefits from a grant from ‘Fondation de France’ (00081132). We thank Life Science Editors for editing assistance.

## Author contributions

EC and IL performed most of the experiments, with the help of CA for metabolic studies, LC for lipophagyi experiments, MM and SLG for AT collection, and SDauv for confocal and videomicroscopy experiments. MD and TM performed the proteomic studies under the supervision of OBS. EC, IL, LN and CM analyzed the data with the help of SDal, PV, and OBS. EC, IL and LN conceived the idea for this project and wrote the manuscript with significant inputs from all authors. LN supervised the study.

## Conflict of interest

The authors declare that no conflict of interest exists.

## Supplementary figure legends

**Supplementary Figure 1:**
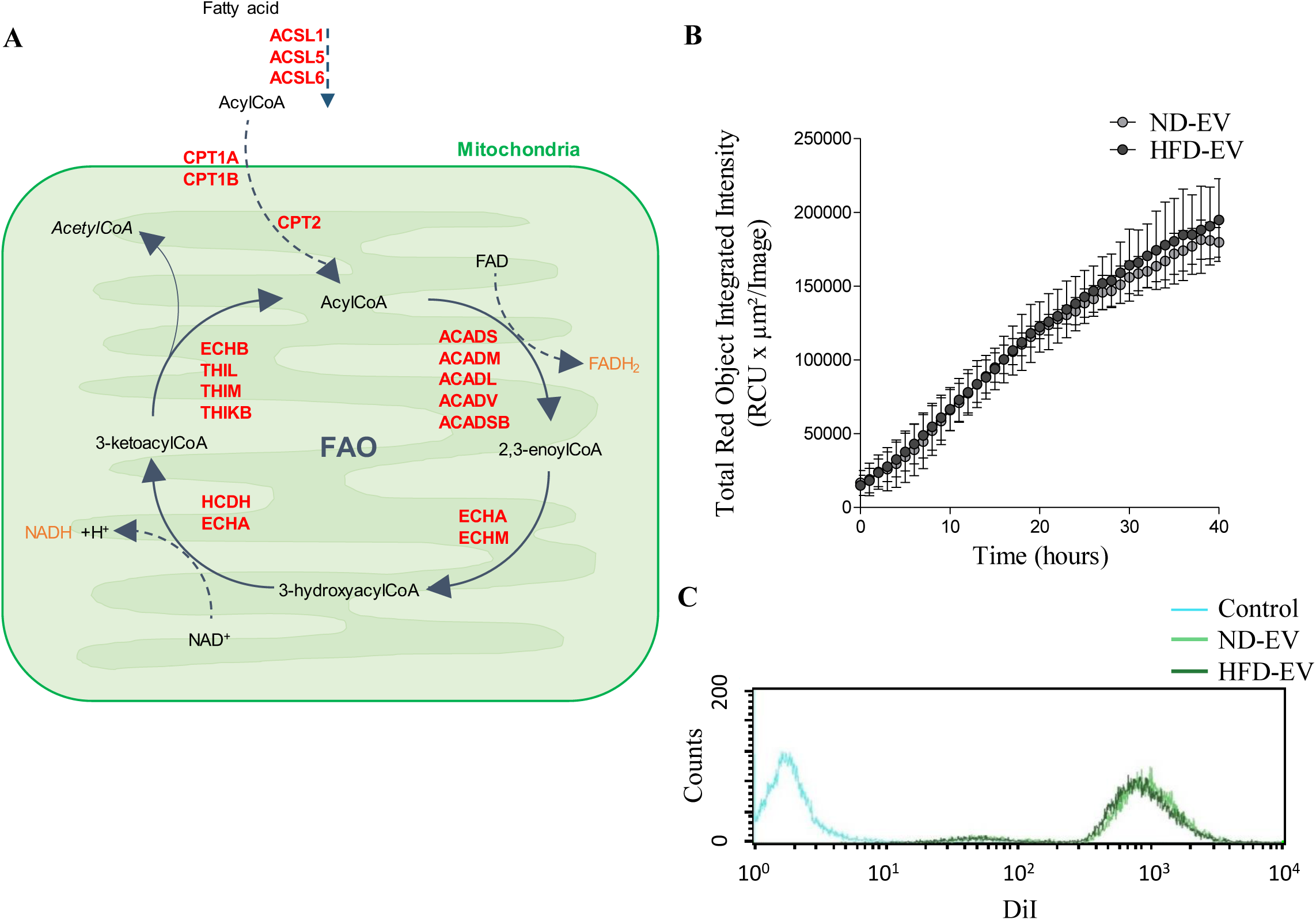
FAO proteins are not upregulated in obese adipocyte EV, and these EV are not preferentially taken up by melanoma cells. A) KEGG pathway analysis using DAVID v6.7 database (http://david.abcc.ncifcrf.gov/) of the proteins that are significantly neither up-, nor downregulated in EV from obese adipocytes revealed a strong association with the FA degradation pathway [p-value 7.4×10^-9^ and corrected (Benjamini-Hochberg) 1.7×10^-9^]. Such proteins identified in equal abundance in obese and lean adipocyte EV are indicated in red on the diagram. B-C) Adipocyte EV from lean (ND) and obese (HFD) mice were fluorescently labeled using the lipophilic DiI probe and added to SKMEL28 cells. B) EV accumulation was evaluated using an Incucyte Zoom system. C) After 48h, cells were harvested and fluorescence was evaluated by flow cytometry (a representative graph is shown).

**Supplementary Figure 2:**
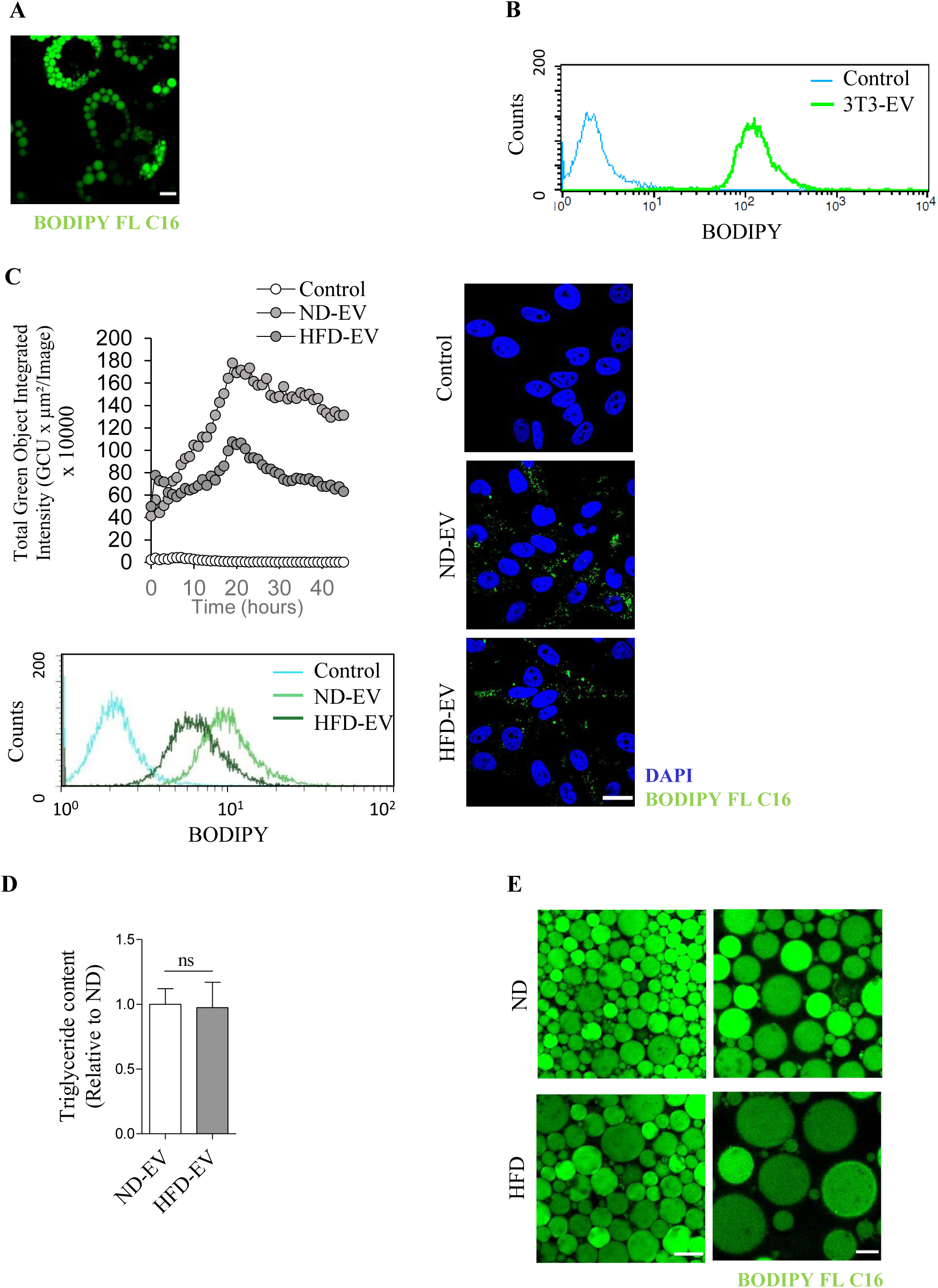
Adipocytes accumulate fluorescent palmitate that is transferred to melanoma cells by EV. A) 3T3-F442A mature adipocytes were loaded with BODIPY FL C16 then fixed and observed by confocal microscopy. Scale bar: 20µm B) SKMEL28 cells exposed to EV secreted by 3T3-F442A adipocytes loaded with BODIPY FL C16 were analyzed by flow cytometry (a representative graph is shown). C) SKMEL28 cells exposed to EV secreted by lean (ND) and obese (HFD) adipocytes previously loaded with BODIPY FL C16. Top left panel, cell fluorescence was assessed using an Incucyte Zoom. Bottom left and right panel, after 24h incubation, cells were harvested and fluorescence was analyzed by flow cytometry (a representative graph is shown) (left) or were fixed and counterstained with DAPI for confocal microscopy observation (right). Scale bar: 20µm. D) Lipids were extracted from EV secreted by adipocytes from lean (ND) and obese (HFD) mice (n=4) and triglyceride content was measured. Bars and error flags represent means ± SEM; statistically significant by unpaired Student’s t-test, ns: non-significant. E) Lean (ND) and obese (HFD) murine adipocytes were loaded with BODIPY FL C16 and observed using confocal microscopy. Scale bars: 100µm (left) or 40µm (right).

**Supplementary Figure 3:**
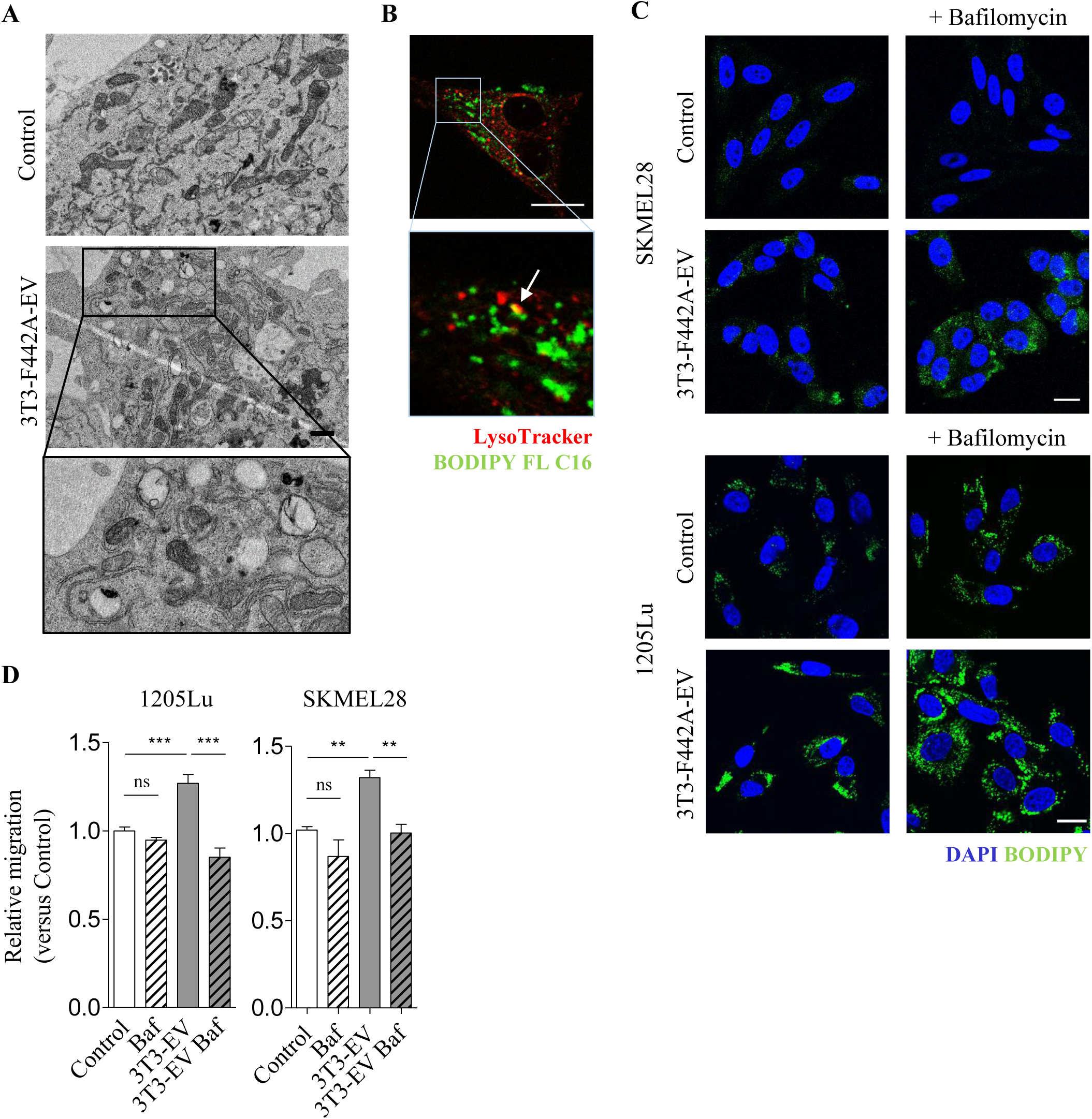
Autophagy is required for lipid degradation and cell migration in response to adipocyte EV. A) Transmission electron micrographs of SKMEL28 cells exposed, or not, to 3T3-F442A EV. A zoomed crop of the area with autophagic structures containing lipids is shown. Scale bar: 1µm. B) SKMEL28 cells were incubated with EV secreted by 3T3-F442A adipocytes previously loaded with BODIPY FL C16. Then, cells were washed, stained with the LysoTracker probe and live cells were observed by confocal microscopy. Scale bar: 20µm. C) Indicated melanoma cells were exposed to 3T3-F442A EV and treated, or not, with Bafilomycin A1. Then, cells were fixed, stained with BODIPY and counterstained with DAPI for observation by confocal microscopy. Scale bar: 20µm. D) SKMEL28 and 1205Lu cells were exposed to 3T3-F442A EV (3T3-EV) and treated, or not, with Bafilomycin A1 (Baf). Cell migration was then evaluated in Boyden chamber assays. Bars and error flags represent means ± SEM (n ≥ 5); statistically significant by one-way ANOVA with post hoc Tukey’s test, *P < 0.05, **P < 0.01, ns: non-significant.

**Supplementary Figure 4:**
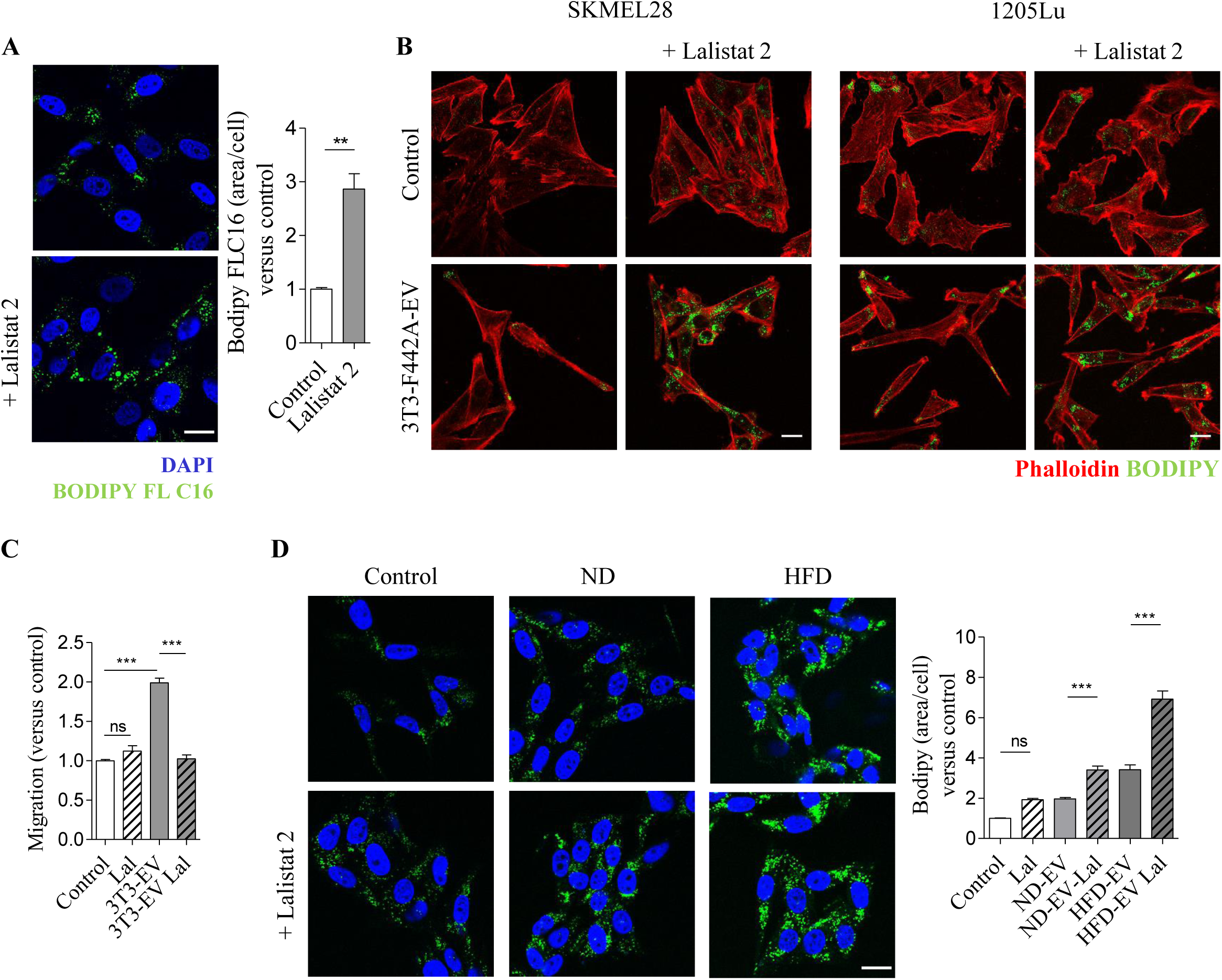
FA transferred from adipocytes to SKMEL28 cells by EV are released from lipid droplets by lipophagy. A) SKMEL28 were incubated with EV secreted by 3T3-F442A adipocytes previously loaded with BODIPY FL C16 and treated, or not, with Lalistat 2. Then, cells were fixed and counterstained with DAPI. Quantification of BODIPY FL C16 staining per cell is shown on the right (n=3). B) SKMEL28 and 1205Lu cells were exposed to 3T3-F442A EV in the presence, or not, of Lalistat 2. Cells were then fixed and stained with BODIPY and Phalloidin. C) SKMEL28 cells were exposed to 3T3-F442A EV (3T3-EV) and treated, or not, with Lalistat 2 (Lal). Cell migration was then evaluated in Boyden chamber assays. D) SKMEL28 cells were exposed, or not, to adipocyte EV from lean (ND) or obese (HFD) mice with, or without, Lalistat 2 (Lal). Cells were then fixed, stained with BODIPY and counterstain with DAPI. Quantification of BODIPY staining per cell is shown on the right (n=5). Scale bars: 20µm. Bars and error flags represent means ± SEM; statistically significant by Student’s t-test (A) or by one-way ANOVA with post hoc Tukey’s test (C-D), **P < 0.005, ***P < 0.001, ns: non-significant.

**Supplementary Figure 5:**
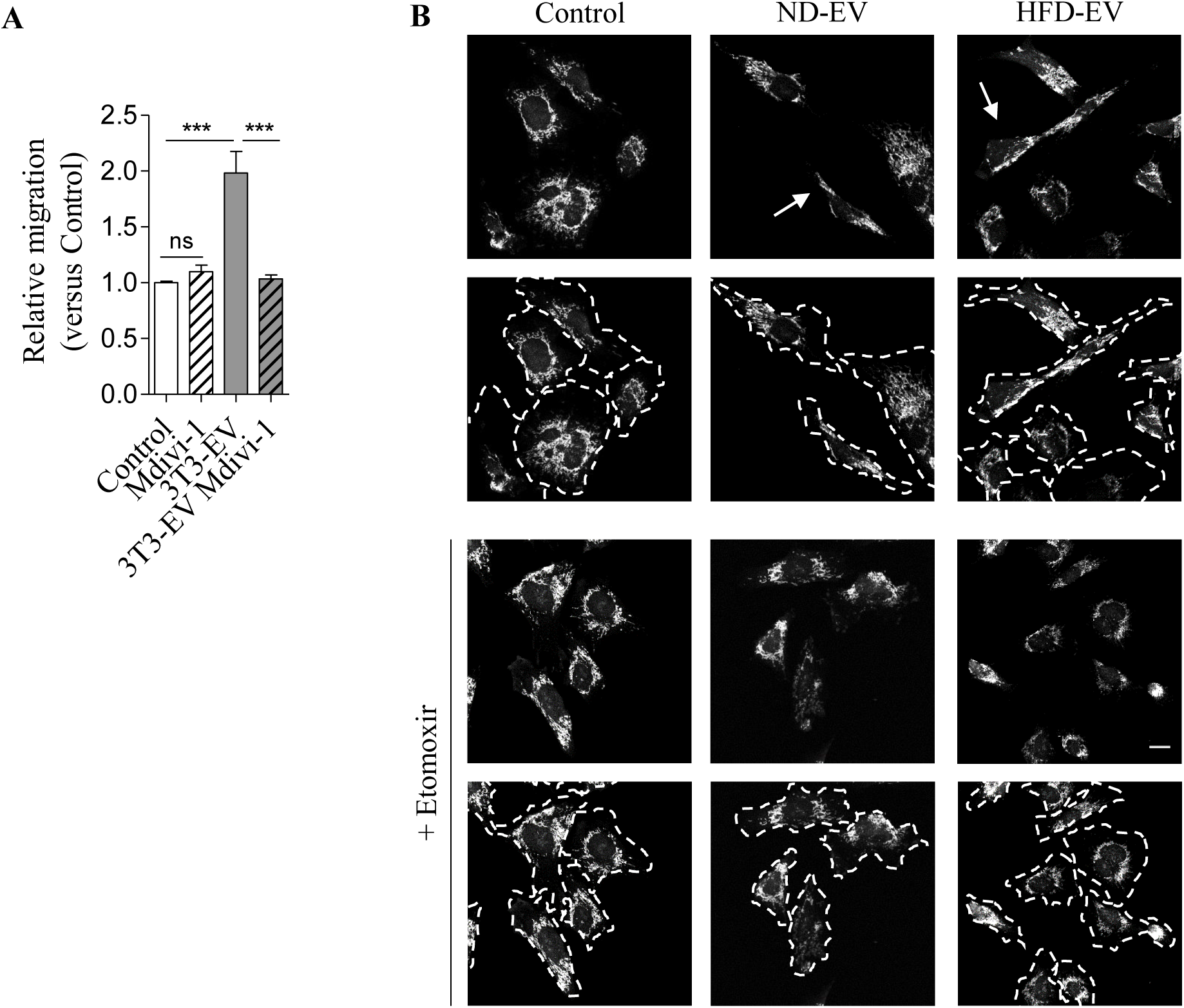
Adipocyte EV modify melanoma cell mitochondrial dynamics, a process that is dependent on FAO and promotes aggressiveness. A) SKMEL28 cells were exposed to 3T3-F442A EV (3T3-EV) and treated, or not, with Mdivi-1. Cell migration was then evaluated in Boyden chamber assays. Bars and error flags represent means ± SEM (n=6); statistically significant determined by one-way ANOVA with post hoc Tukey’s test, ***P<0.001, ns: non-significant. B) 1205Lu cells were exposed to adipocyte EV from lean (ND) or obese (HFD) mice for 48 with, or without, Etomoxir for the last 24h. Cells were then stained with a MitoTracker probe, fixed and observed by confocal microscopy. Arrows indicate the presence of mitochondria in membrane protrusions. Underneath each original image, an image depicting the outline of cells in dotted lines is shown. Scale bar: 20µm.

**Supplementary Figure 6:**
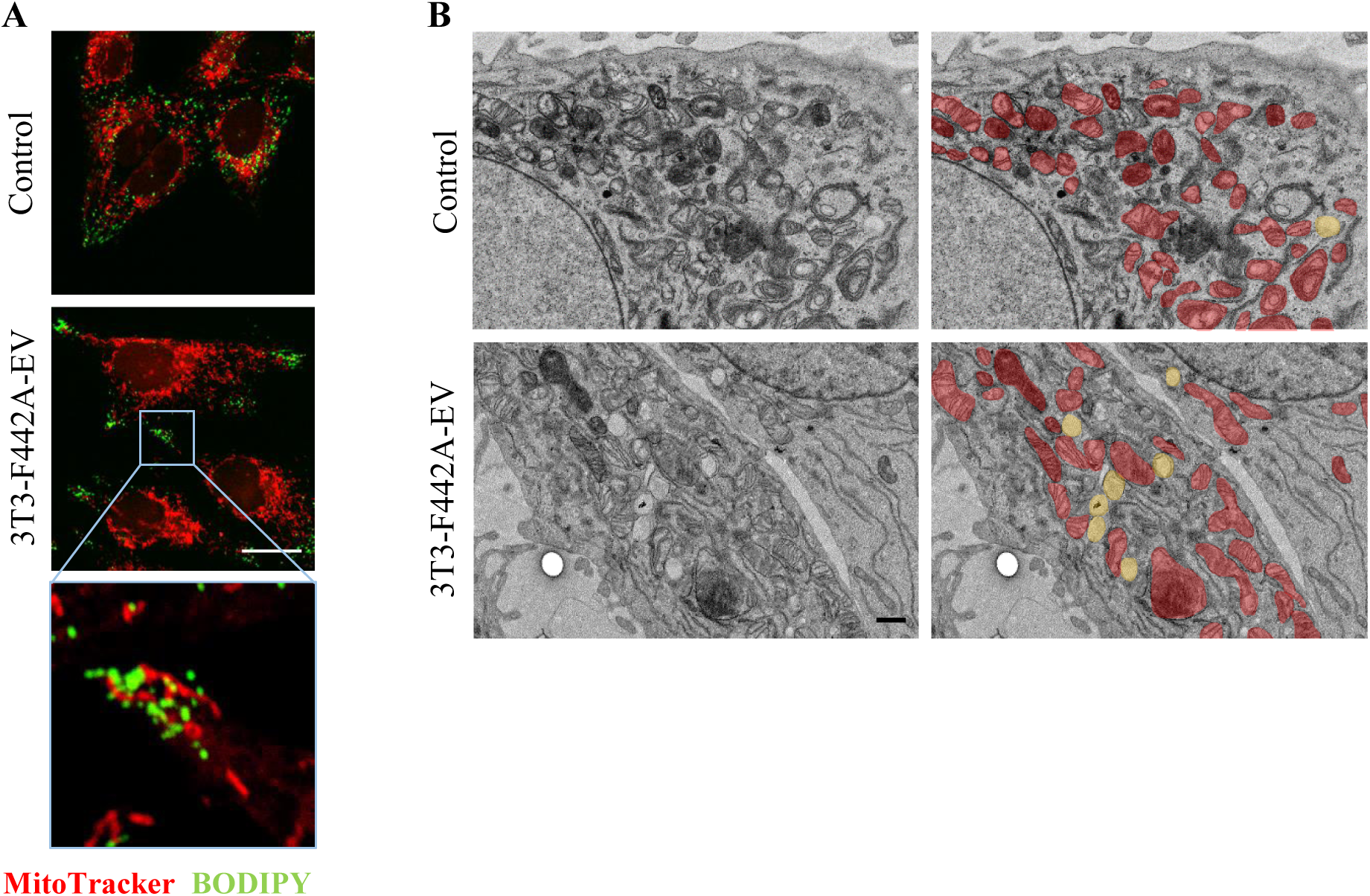
Lipid droplets are found in membrane protusions, at proximity to mitochondria, in melanoma SKMEL28 cells exposed to adipocyte EV. A) SKMEL28 cells exposed to 3T3-F442A EV were stained with a MitoTracker probe. Then, cells were fixed, stained with BODIPY and observed by confocal microscopy. A zoomed crop of the area containing mitochondria and lipid droplets in a membrane protrusion is shown. B) Transmission electron microscope observations of SKMEL28 cells exposed, or not, to 3T3-F442A EV. Mitochondria are colored in red and lipid droplets are colored in yellow on images on the right. Scale bars represent 20µm for confocal microscopy images and 1µm for electron microscopy images.

